# The Hippo/YAP Pathway Mediates the De-differentiation of Corneal Epithelial Cells into Functional Limbal Epithelial Stem Cells *In Vivo*

**DOI:** 10.1101/2024.06.11.596348

**Authors:** Yijian Li, Lingling Ge, Bangqi Ren, Xue Zhang, Zhiyuan Yin, Hongling Liu, Yuli Yang, Yong Liu, Haiwei Xu

## Abstract

Regeneration after tissues injury is often associated with cell fate plasticity, which restores damaged or lost cells. The de-differentiation of corneal epithelial cells (CECs) into functional stem cells after the ablation of innate stem cells, known as limbal epithelial stem cells (LESCs), remains controversial. In this study, we showed the functional maintenance of corneal epithelium after the ablation of innate stem cells, and the regeneration of functional LESCs, which maintained corneal transparency, prevented corneal conjunctivalization and participated in the wound healing. Subsequent intravital lineage tracing revealed that CECs could de-differentiate into active or quiescent LESCs, which functioned as well as their innate counterparts. Furthermore, the de-differentiation of CECs required an intact limbal niche, and the outcome of the competition between conjunctival and corneal epithelium for the limbal niche determined whether the de-differentiation would occur or not. Mechanically, the suppression of YAP signal promoted the de-differentiation of CECs after the ablation of innate stem cells, while the persistent activation of YAP prevented the de-differentiation of CECs after an additional alkali burn to the limbal stroma. These results will pave the way for an alternative approach to treat limbal stem cell deficiency (LSCD) by modulating the de-differentiation of CECs *in vivo*.

## 1. Introduction

The limbus, where limbal epithelial stem cells (LESCs) reside exclusively in its basal layer, is a narrow ring-shaped transition zone between cornea and conjunctiva (1–3). The zone forms a boundary to prevent conjunctival epithelium from invading the corneal surface. LESCs are responsible for the continuous renewal of corneal epithelium by producing transit-amplifying cells (TACs), which divide, migrate centripetally toward the central cornea, differentiate into mature corneal epithelial cells (CECs) and eventually desquamate (4). LESCs exhibit remarkable proliferative capacity, whereas TACs are more limited in this capacity, demonstrating a hierarchy of proliferative capacity (5). Due to the existence of TACs, corneal epithelium can self-maintain and repair small central corneal wound independent of LESCs (6–8). Only after extensive corneal epithelial wound, LESCs rapidly proliferate to compensate for the loss of CECs (8–10). Recent findings have revealed the co-existence of two LESCs sub-populations localized in distinct and well-defined sub-compartments in mice: active LESCs (aLESCs) in the inner limbus undergo mostly symmetric divisions and are required to sustain the population of TACs that support corneal homeostasis; quiescent LESCs (qLESCs) in the outer limbus participate in the boundary formation and wound healing after large wound (11, 12). In addition, as the fast-renewal of cornea, LESCs are found to be abundant and equipotent, and follow the stochastic competition model of stem cells, where equipotent stem cells neutrally compete with each other for the niche of stem cells. The neutral competition between stem cells at the population level will lead to a stochastic pattern of behavior known as neutral drift dynamics, where the tissue is maintained by an ever-diminishing clone number of ever-increasing size (11, 13, 14).

Given the crucial roles of LESCs in the cornea, any pathology which directly damages them or disrupts the limbal niche, a specialized microenvironment of LESCs, can potentially cause limbal stem cell deficiency (LSCD). LSCD is a rare, progressive ocular surface disorder which leads to corneal conjunctivalization, neovascularization and opacification (15–17). Consequently, it is generally believed that the loss of LESCs can’t be repaired and unavoidably results in blindness. Interestingly, the ablation of the entire limbal epithelial cells (including LESCs) won’t result in any symptoms of LSCD (18, 19). One study reported that corneal-committed cells could migrate back to limbus, de-differentiate into CK15-GFP^+^ stem cells and reform the tissue boundary (20). However, recent studies have refuted both the phenomenon of de-differentiation and even the centrifugal migration of CECs after the removal of limbal epithelium (12, 21). These contradictory conclusions are never resolved.

The highly conserved Hippo pathway plays roles in development, regeneration and the progression of several diseases. The canonical Hippo cascade is initiated by MST1/2, which activates LATS1/2 through phosphorylation. Activated LATS1/2 then phosphorylates (e.g., YAP Ser127) and inactivates YAP/TAZ by repressing their nuclear translocation. YAP/TAZ functions as transcriptional co-factor and interacts with transcriptional factors (e.g. TEAD, p63, and AP1) to regulate a large number of genes, which are involved in cell proliferation, plasticity and fate determination that are essential for tissue regeneration and repair (22, 23). In the corneal epithelium, YAP has been shown to participate in the maintenance of corneal epithelial progenitor cells and wound healing (10, 24). Previous studies also indicated that transient activation of YAP could locally induce a hyper-proliferative state of CECs with increased expression of CK14 and p63α (potential LESCs markers) (10, 25). Whether the Hippo/YAP pathway plays roles in the de-differentiation of CECs needs further exploration.

Here, we demonstrated that CECs were able to de-differentiate into functional LESCs, including both aLESCs and qLESCs, thereby regenerating the LESCs pool and restoring functions of the limbus. Additionally, we revealed the competition between corneal and conjunctival epithelial cells for the limbal niche and the outcome of this competition determined the de-differentiation of CECs occurred or not, which reconciled the apparent contradiction in this field. Furthermore, the interpretation of the role of Hippo/YAP pathway in the de-differentiation of CECs may pave the way for an alternative and efficient treatment of LSCD by modulating the fate of CECs.

## 2. Materials and methods

### 2.1 Animals, lineage tracing and intravital imaging

Animal care and use conformed to the guidelines of the Laboratory Animal Welfare and Ethics Committee at the Third Military Medical University (Army Medical University), Chongqing, China. The ethics approval number is AMUWEC20197030. All mice and Long Evans rats were housed under specific-pathogen-free (SPF) conditions. CK14-CreERT2 (Shanghai Model Organisms; NM-KI-190024), Slc1a3-CreERT (Jackson Laboratory; 012586) and reporter mice (GemPharmatech; H11-loxP-ZsGreen-stop-loxP-tdTomato; T057726) were purchased and crossed to generate double-transgenic mice (CK14/Slc1a3-CreERT; H11-reporter). The genotypes of mice were identified by PCR analysis of tail DNA according to the manufacturer’s instructions (Mouse Direct PCR Kit, Bimake, B40013). Heterozygous double-transgenic mice aged 4-6 months were used for lineage tracing. For lineage tracing experiments, Cre recombinase activity was induced by injecting intraperitoneally (200μL) 2mg/day Tamoxifen (TargetMol, T6906) dissolved in corn oil for five consecutive days. These tdTomato-labeled LESCs or TACs would then proliferate and migrate centripetally toward the central cornea, resulting in the formation of stripes. For intravital imaging of the eyes, mice were initially anesthetized with 2.5% isoflurane (RWD Life Science), and the head was stabilized with the eye to be imaged under an Olympus BX51 fluorescence microscope. Animals were returned to the housing facility after woke up.

### 2.2 LESCs ablation, cornea epithelial wound healing, NaOH application and tLSCD model

For innate LESCs ablation, Long Evans rats and C57BL/6J or lineage-tracing mice aged 6 months were anesthetized by 2.5% isoflurane. After application of topical proparacaine (0.5%), 2mm-wide limbal epithelium (including surrounding corneal and conjunctival epithelium) was scratched off using a tunnel knife under a dissecting microscope. To investigate the competition between corneal and conjunctival epithelium for the limbus, different-sized limbal and corneal epithelium scratch-off was performed as depicted in Fig. 5E. For corneal epithelial wound and healing, small (1mm-diameter), medium (1.5mm-diameter) or large (2mm-diameter) wound of mice and 3.5mm-diameter wound of rats were performed according to the experimental design. For the localized application of NaOH after the innate LESCs ablation, a 1mm-wide filter paper strip was soaked in 1M NaOH for 5s, picked up in the air for 5s, and applied on the temporal limbus (1/4 - 1/5 of the whole limbus) for 5s followed by extensive washing with water. For the total LSCD model (tLSCD), a two-step scratch-off of limbal and corneal epithelium was performed as depicted in Fig. S5B. In brief, limbal epithelium was scratched off and stained by fluorescein sodium to make sure the ablation of LESCs, then corneal epithelium was scratched off. All these defects were visualized and imaged by 1% fluorescein sodium staining to confirm successful removal of epithelium and subsequent examination of wound healing.

### 2.3 Drug treatment

All mice were treated with local ocular administration for four consecutive days, and their corneas were collected at 6 days after limbal epithelial removal for subsequent experiments. Vehicle (0.2% DMSO), verteporfin (VTP, 20μM, Selleck, S1786), or TRULI (20μM, CSNpharm, CSN26140) was administrated to ocular surface every 8 hours after LESCs ablation either with or without NaOH application. Manual blinking was performed to make sure complete absorption of drug solutions by ocular surface. In experiments involving VTP, mice were housed in darkness to protect them from direct sunlight.

### 2.4 Fluorescence staining

For frozen sections, corneal tissues of mice and rats were fixed in 4% paraformaldehyde (PFA) for 30 minutes, washed with PBS, incubated in 30% sucrose for 4 hours and embedded in Tissue-Tek OCT compound. Frozen sections of corneal tissues (14μm thick) were cut using a Leica cryostat, and mounted on microscope slides. Tissue sections were permeabilized (0.3% Triton X-100 for 10 minutes) and blocked (3% BSA for 60 minutes). Following these treatments, tissue sections were incubated with primary antibodies (16 hours, 4°C), secondary antibodies (60 minutes, 37°C), DAPI and mounted with anti-fade medium. For whole-mount cornea staining, corneas were isolated with iris removal, fixed in 4% PFA for 1 hour, permeabilized and blocked (1% Triton X-100 and 3% BSA, 4 hours) and incubated with primary antibodies (1% Triton X-100 and 3% BSA, 24 hours, 4°C on a shaker). After washes with 0.3% Triton X-100, corneas were incubated with secondary antibodies (1% Triton X-100 and 3% BSA, 16 hours, 20°C on a shaker), followed by DAPI staining, tissue flattening under a dissecting binocular and mounting. The limbus was localized as the junction between iris and ciliary body.

The following primary antibodies were used: rabbit anti-CK12 monoclonal antibody (Abcam, ab185627; 1:400), rabbit anti-CK7 monoclonal antibody (Abcam, ab181598; 1:400), rabbit anti-connexin 43 (Cx43) monoclonal antibody (CST, #3512; 1:80), rabbit anti-ApoE monoclonal antibody (Abcam, ab183596; 1:400), rabbit anti-CK14 monoclonal antibody (Abcam, ab119695; 1:200), rabbit anti-deltaN-p63 polyclonal antibody (BioLegend, 619002; 1:400), rabbit anti-p75NTR monoclonal antibody (CST, #8238; 1:800), rabbit anti-CD63 polyclonal antibody (Bioworld, BS72936; 1:400), rabbit anti-TSPAN7 polyclonal antibody (Proteintech, 18695-1-AP; 1:100), rabbit anti-IFITM3 monoclonal antibody (CST, #59212; 1:100), mouse anti-ATF3 monoclonal antibody (Santa Cruz, sc-518032; 1:100), rabbit anti-YAP monoclonal antibody (CST, #14074; 1:100), rabbit anti-p-YAP-Ser127 monoclonal antibody (Abcam, ab76252; 1:100), rabbit anti-active-YAP (non-phosphorylated YAP) monoclonal antibody (Abcam, ab205270; 1:200), rabbit anti-Ki67 monoclonal antibody (CST, #9129; 1:100), rabbit anti-p-Histone H3-Ser10 (pH3) monoclonal antibody (CST, #53348; 1:100) and mouse anti-CK15 monoclonal antibody (Santa Cruz, sc-47697; 1:200). The following secondary antibodies were used: Goat anti-rabbit Alexa-Fluor-488 (Invitrogen, A32731TR; 1:400), Goat anti-mouse Alexa-Fluor-488 (Invitrogen, A11001; 1:400), Goat anti-rabbit Alexa-Fluor-647 (Invitrogen, A32733; 1:600) and Goat anti-rabbit Alexa-Fluor-568 (Invitrogen, A11011; 1:400).

For EdU labeling and staining, single intraperitoneal injection of mice at indicated time point with 200uL (6mg/mL) EdU (Beyotime) was performed, and mice were euthanized 4 hours later with inhalational CO_2_. Corneas were fixed in 4% PFA for 1 hour, and stained according to the manufacturer’s instructions (BeyoClickiTM, Beyotime) after whole-mount staining protocol. For F-actin staining, whole-mount cornea or tissue sections were incubated in fluorescein-phalloidin (Invitrogen, A12380; 1:800; cornea for 30 minutes, sections for 15minutes) followed by DAPI staining.

Images were acquired by Olympus BX51 fluorescence microscope and Zeiss LSM800 confocal microscope, and processed by Image J software.

### 2.5 Quantification and statistical analysis

For the quantification of the mean fluorescence intensity (MFI), all experimental conditions and acquisition parameters of confocal microscope were identical, and these regions that were determined were manually drawn and MFI was quantified by Image J software. The percentage of ApoE^+^ cells at the limbus, and MFI(limbal basal cells)/MFI(peripheral basal cells) of Cx43 and CK12 were determined to be indexes of the recovery of LESCs. For the quantification of cellular area, the boundary of cell was manually drawn based on the F-actin staining, and cellular area was quantified by Image J software.

Statistical analysis was conducted using one-way ANOVA with Tukey’s test for multi-group comparisons, and Student’s t-test for comparison between two groups. Mean ± SD were represented, and significance was set as *p < 0.05, **p < 0.01, ***p < 0.001 and ****p < 0.0001, unless otherwise indicated. The sample sizes (n) were indicated in the figure legends. All analysis were performed with GraphPad Prism software (Version 6.0).

## 3. Results

### 3.1 Functional Maintenance of Corneal Epithelium and Regeneration of LESCs after the Ablation of Innate Stem Cells

We ablated innate stem cells (i.e., LESCs) of cornea in mice by scratching off an approximately 2mm-wide ring-shaped zone, including the entire limbal epithelium, marginal conjunctival and corneal epithelium. Fluorescein sodium staining showed that this limbal wound could be healed within 24 hours, and the cornea sustained transparency for up to 6 months (Fig. 1B). Immunostaining for CK12 (a marker of CECs) showed the maintenance of CECs across the whole cornea (Fig. 1B), and CK7 (a marker of conjunctival epithelial cells) immunostaining showed that no conjunctival epithelial cells invaded the cornea (Fig. 1B; Fig. S1A). This implied the functional maintenance of corneal epithelium during homeostasis after the ablation of innate LESCs.

**Figure 1:**
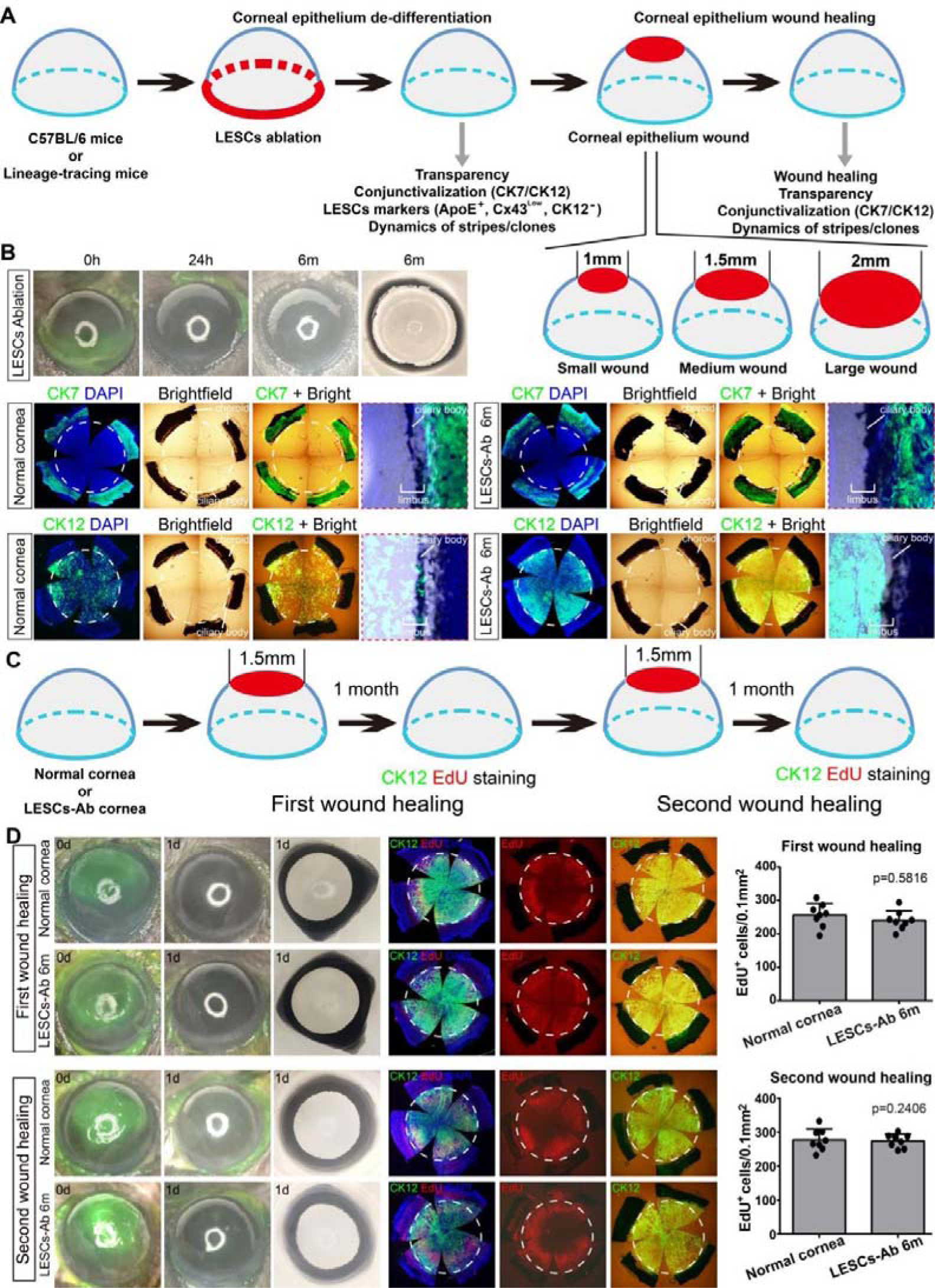
The functional maintenance of corneal epithelium after innate stem cells ablation. (**A**) Experimental strategy to resolve the regeneration of LESCs after innate stem cells ablation *in vivo*. The transprarency, conjunctivalization and two consecutive central corneal wound healing are shown in Figure 1. The expressions of LESCs markers and dynamics of stripes/clones are shown in Figure 2 and Figure 3-4, respectively. The different-sized wounds (small, medium and large) of central corneal epithelium in mice (Figure 4) are also shown. (**B**) The limbal epithelium and marginal corneal and conjunctival epithelium of 6-month-old C56BL/6J mice were removed by surgery. Fluorescein sodium staining and bright-field images of injured eyes at indicated days are shown. Whole-mount normal and LESCs-ablation (LESCs-Ab; 6 months) corneas were immunostained for conjunctival epithelial marker CK7 and corneal epithelial marker CK12. (**C**) Experimental strategy of two consecutive central corneal wound healing of normal and LESCs-ablation corneas. (**D**) Fluorescein sodium images and whole-mount staining of normal and LESCs-ablation corneas for EdU (cell proliferation) and CK12 after the first and second wound healing. Dashed circles indicate the limbus, which is localized based on the location of iris and ciliary body in the bright field. These EdU-positive proliferative cells at 8 zones of peripheral cornea are quantified. Data are the mean ± SD; statistical analysis are performed by Student’s unpaired t-test, and p values are shown.

Previous studies suggested that corneal epithelium had limited self-renewal and therefore poor regenerative or repairing capacity (6, 19, 21). To explore whether the maintenance of corneal epithelium after innate LESCs ablation resulted from this limited capacity of self-renewal, we performed two consecutive central corneal wound healing on both normal and LESCs-ablation corneas (Fig. 1C; Fig. S1B). During both the first and second central corneal wound healing, both normal and LESCs-ablation corneas repaired the wound and sustained transparency without corneal conjunctivalization (Fig. 1D; Fig. S1C, D). More surprisingly, LESCs-ablation corneas showed identical healing rates and proliferative patterns and abilities (EdU staining) to those of normal corneas during both the first and second wound healing (Fig. 1D). Thus, these results ruled out the possibility that this functional maintenance of corneal epithelium was due to the limited self-renewal of CECs, and instead implied the regeneration of LESCs following the ablation of innate LESCs.

As known, a single definitive marker specific for LESCs remains infeasible. Instead, a combination of a set of markers and functional descriptions is required to define LESCs (17, 26, 27). In this context, we tested various potential LESCs markers (including CK14, p63, ApoE, p75NTR, CD63, TSPAN7, IFITM3 and ATF3) based on previous reports and single-cell RNA sequencing (11, 28). Consistent with previous studies, most of these markers showed to be non-specific for LESCs (Fig. 2A). Ultimately, we defined LESCs with the following set of markers: ApoE^+^/Cx43^low^/CK12^-^. To investigate whether the regeneration of LESCs occurred, we detected the expressions of these markers at the limbus after the ablation of innate LESCs. Epithelial cells at the limbus began to express ApoE at 6 days after the ablation of innate LESCs and sustained ApoE expression for at least 6 months (Fig. 2B-E). Concurrently, epithelial cells at the limbus showed low Cx43 expression at 6 days and sustained for at least 6 months (Fig. 2B-E). Intriguingly, the CK12 level of basal epithelial cells at the limbus decreased at 4 days, and some limbal basal epithelial cells didn’t express CK12 at 20 days (Fig. 2B-E). At 20 days after the ablation of innate LESCs, expressions of ApoE, Cx43 and CK12 approximately recovered to their levels during normal homeostasis (Fig. 2E). This functional maintenance of cornea and the regeneration of Cx43^low^/CK12^-^ cells at the limbus were also observed in a rat model of LESCs ablation (Fig. S2). Together with aforementioned functional maintenance of corneal epithelium (cornea transparency, no conjunctivalization and identical wound healing rate), these results strongly indicated the regeneration of LESCs after the ablation of innate stem cells.

**Figure 2:**
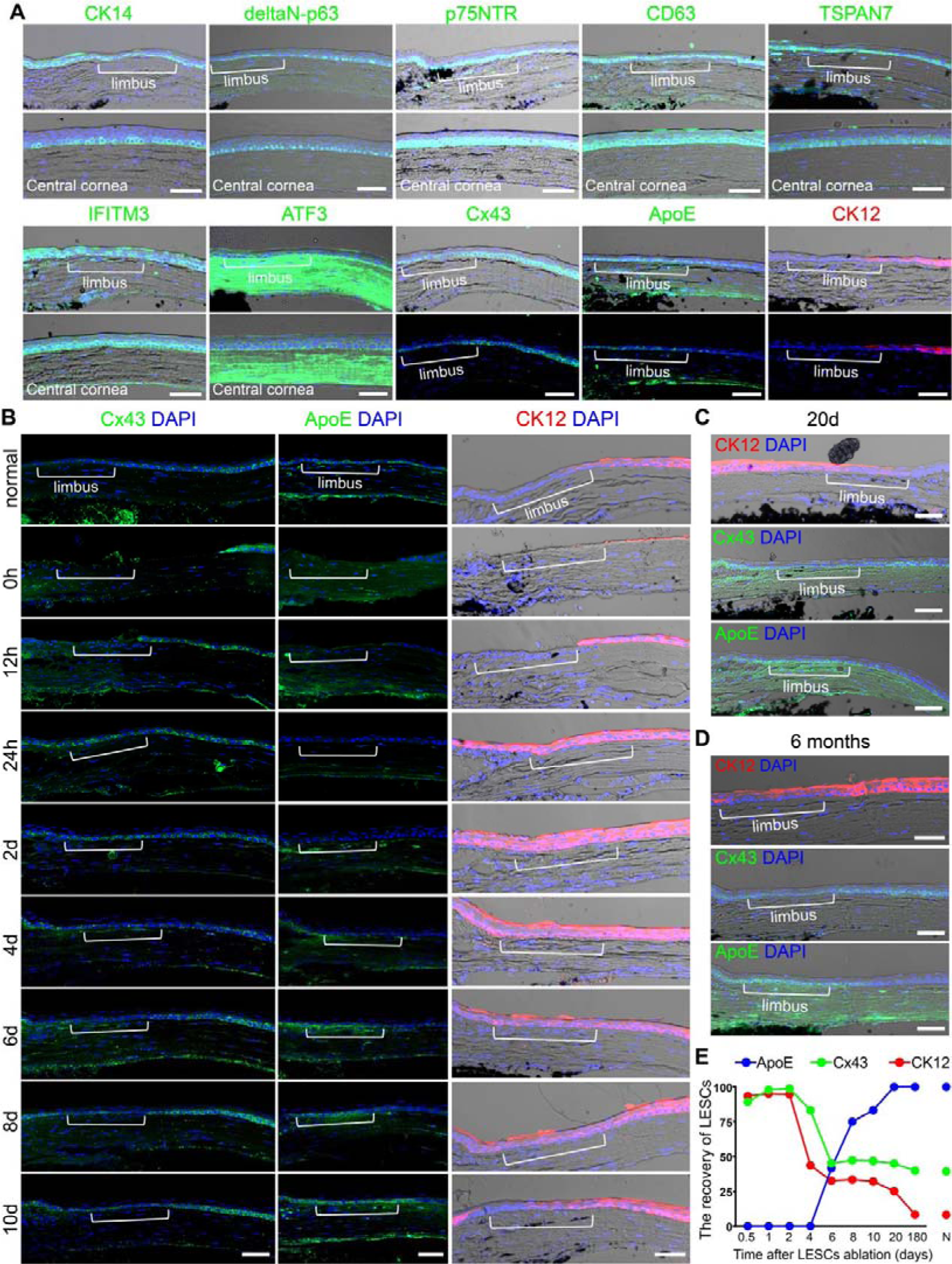
The regeneration of LESCs at limbus after innate stem cells ablation. (**A**) Immunostaining of potential LESCs markers (CK14, deltaN-p63, p75NTR, CD63, TSPAN7, IFITM3, ATF3 and ApoE), Cx43 and CECs marker CK12 in frozen sections of normal cornea. (**B-D**) Immunostaining of Cx43, ApoE and CK12 in frozen sections of normal cornea and LESCs-ablation cornea at indicated days after the limbal epithelium removal. The limbus was shown. (**E**) The percentage of ApoE^+^ cells at the limbus, and MFI(limbal basal cells)/MFI(peripheral basal cells) of Cx43 and CK12 were determined to indicate the recovery of LESCs. N, normal cornea. Scale bars, 50 um. Note that the limbus is localized based on the location of iris and ciliary body in the bright field and the expressions of LESCs markers.

### 3.2 The De-differentiation of CECs into aLESCs *In Vivo*

These regenerated LESCs can derive from 1) residual LESCs, 2) de-differentiation of CECs, and 3) trans-differentiation of conjunctival cells or others. As shown in Fig1B and Fig2B, there are no residual epithelial cells at the limbus, making it highly impossible that residual LESCs mainly contribute to the regeneration of LESCs. Instead, the observed reduction of CK12 expression in basal CECs at the limbus implied the possibility that these regenerated LESCs resulted from the de-differentiation of CECs. To explore this hypothesis, we performed lineage-tracing experiments using heterozygous 6-month-old CK14-CreERT2;H11-reporter mice. Upon transient exposure to tamoxifen, both corneal and conjunctival epithelial cells were labeled, and some tdTomato-labeled LESCs or TACs proliferated and migrated centripetally, resulting in the formation of corneal epithelial stripes (Fig. S3A-C). These aLESCs-derived stripes can extend to the central cornea and are long-lived, while TACs-derived stripes are shorter and only last for less than 4 weeks during homeostasis and wound healing. These qLESCs can only form small cell clones or clusters during homeostasis, and proliferate and migrate toward the central cornea and form stripes during wound healing. Some aLESCs-derived tdTomato^+^ stripes disappeared gradually (Fig. S3C), which is consistent with these features of aLESCs that follow the rule of stochastic competition and neutral drift (11, 13, 14).

Next, we ablated innate LESCs and the dynamics of tdTomato-labeled corneal epithelial stripes were monitored by intravital imaging. The tdTomato-labeled stripe migrated back to the limbus within 2 days, grew longer gradually and reached the central cornea with the stripe “foot” remained stable at the limbus for 10 weeks, which is the feature of aLESCs-derived stripes. Intriguingly, the stripe “foot” detached from the limbus at 15 weeks, and the stripe vanished gradually and became extinct by 24 weeks (Fig. 3A). However, the zone of the disappeared stripe was still CK12^+^ CECs after the healing of a medium central corneal wound (Fig. 3B, C②). These results implied that these tdTomato-labeled cells de-differentiated into aLESCs, but they lost the stochastic competition between tdTomato-unlabeled aLESCs. To observe the dynamics of these de-differentiated aLESCs during central corneal wound healing, we performed a medium wound on cornea when the tdTomato-labeled stripe still had a “foot” at the limbus. The tdTomato-labeled stripe proliferated, migrated centripetally toward the central cornea, and were CK12^+^ CECs (Fig. 3D-F①). Confusingly, not all zones of the cornea of CK14-lineage-tracing mice were CK12^+^ corneal epithelium (Fig. 3C, F). This might result from the destruction of endogenous CK14 gene by the insertion of CreERT2, which might affect CECs’ characteristic and their capability of de-differentiation (Fig. S3A, D). Taken together, these results suggested the de-differentiation of CECs into aLESCs after the ablation of innate stem cells, and these aLESCs participated in corneal epithelial homeostasis and wound healing.

**Figure 3:**
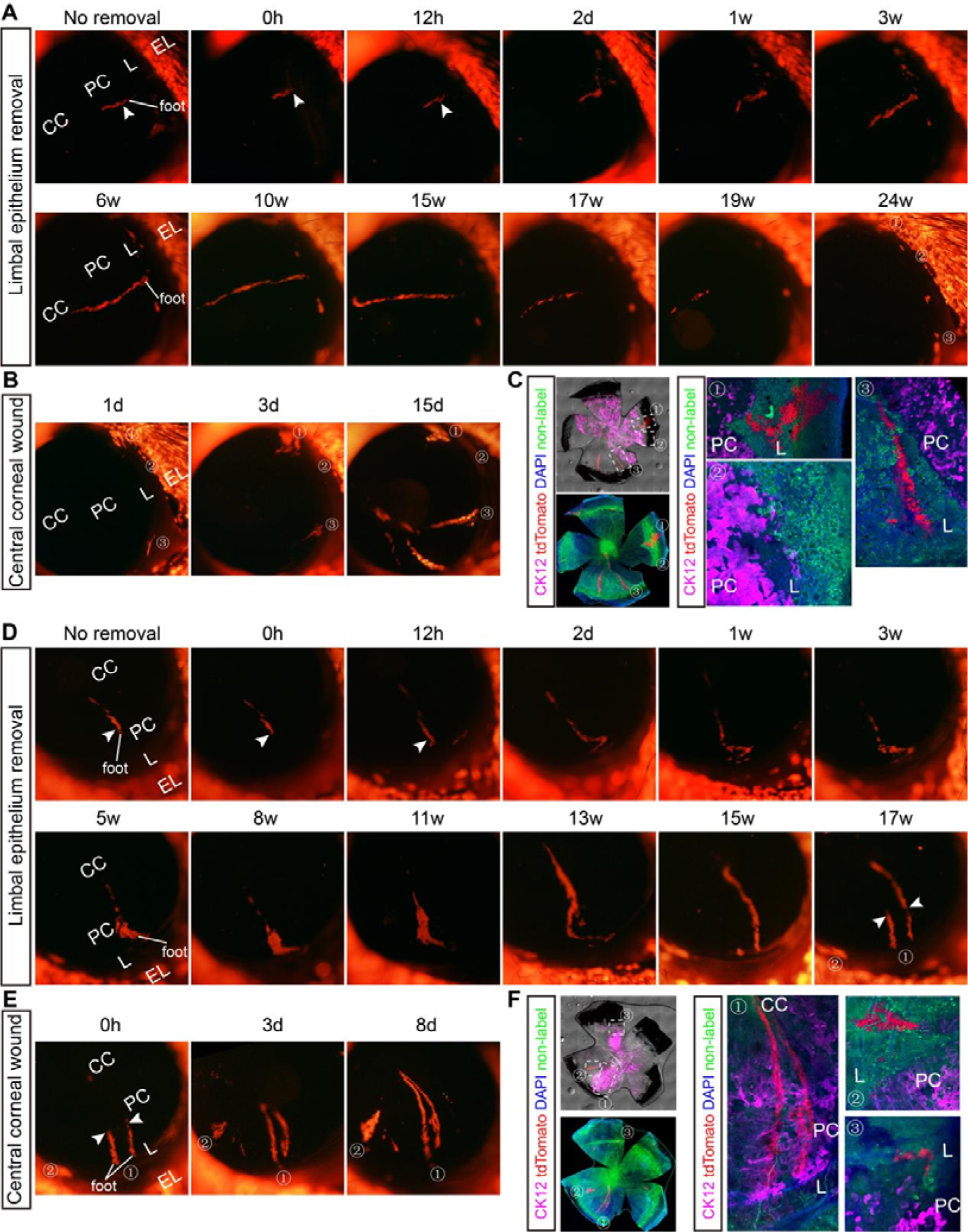
The de-differentiation of CECs into aLESCs *in vivo*. (**A-C**) An example of de-differentiated aLESCs that lose the stochastic competition between stem cells. (**A**) The 6-month-old transgenic mice (CK14-CreERT2; H11-reporter) were induced to express tdTomato reporter and limbal epithelial removal (white arrowheads) was performed after the formation of tdTomato^+^ stripe. Intravital tracking of accurate stripe was followed by fluorescent microscopy over time. (**B**) The medium wound of central corneal epithelium was performed at 24 weeks after limbal epithelial removal, and the changes of lineage-tracing tdTomato^+^ cells were investigated during the wound healing. (**C**) The whole-mount CK12 immunostaining of lineage-tracing corneas at 15 days after the wound of central corneal epithelium. (**D-F**) An example of de-differentiated aLESCs that maintain the stripe. (**D**) Limbal epithelial removal (white arrowheads) was performed after the formation of tdTomato^+^ stripe. Intravital tracking of accurate stripe was followed by fluorescent microscopy over time. (**E**) The medium wound of central corneal epithelium was performed (white arrowheads) at 17 weeks after limbal epithelial removal, and the changes of lineage-tracing tdTomato^+^ cells were investigated during the wound healing. (**F**) The whole-mount CK12 immunostaining of lineage-tracing corneas at 8 days after the wound of central corneal epithelium. The limbus is localized based on the location of iris and ciliary body in the bright field of the whole-mount cornea. The “foot” means the end of corneal epithelial stripe that is closer to the limbus. EL, eyelid; L, limbus; PC, peripheral cornea; CC, central cornea.

### 3.3 The De-differentiation of CECs into qLESCs *In Vivo*

Interestingly, nearly all tdTomato-labeled corneal epithelial stripes of CK14-lineage-tracing mice de-differentiated into aLESCs, leading us to investigate whether CECs could also de-differentiate into qLESCs. We used Slc1a3-CreERT;H11-reporter mice, which were used to explore the compartmentalization of stem cell populations across the ocular surface in a previous report (29), to induce stripes, performed limbal epithelial removal and monitored by intravital imaging. These tdTomato-labeled stripes migrated back to the limbus, disappeared gradually, became small clones at about 3 weeks and remained at the limbus during the whole experiments (Fig. 4A, D; Fig. S4A).

**Figure 4:**
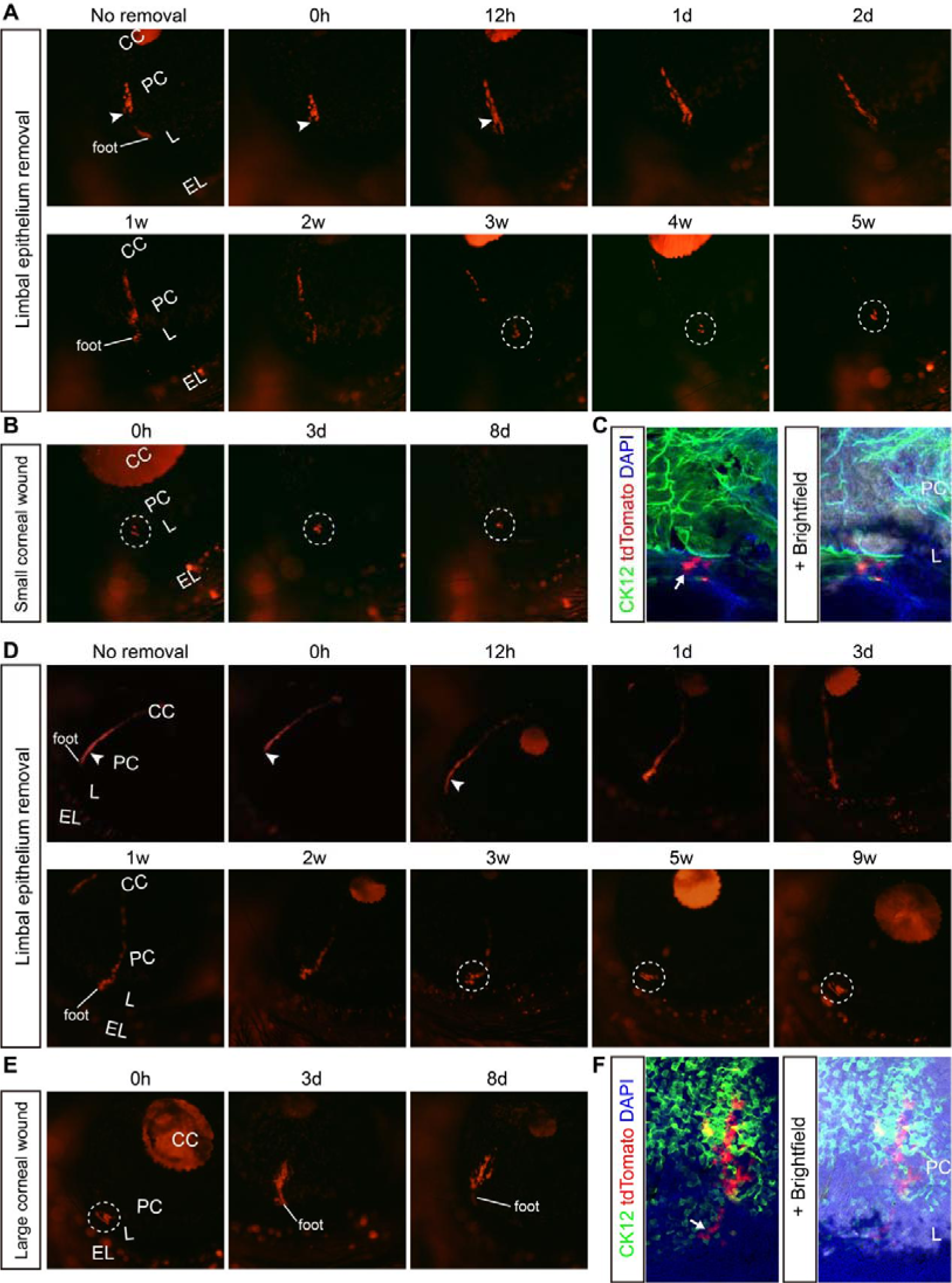
The de-differentiation of CECs into qLESCs *in vivo*. (**A-C**) The response of de-differentiated qLESCs to small wound of central corneal epithelium. (**A**) The 6-month-old transgenic mice (Slc1a3-CreERT; H11-reporter) were induced to express tdTomato reporter and limbal epithelial removal (white arrowhead) was performed after the formation of tdTomato^+^ stripe. Intravital tracking of accurate stripe was followed by fluorescent microscopy over time. (**B**) The small wound of central corneal epithelium was performed at 5 weeks after limbal epithelial removal, and the change of lineage-tracing tdTomato^+^ clone was investigated during the wound healing. (**C**) The whole-mount CK12 immunostaining of lineage-tracing corneas at 8 days after the small wound of central corneal epithelium. (**D-F**) The response of de-differentiated qLESCs to large wound of central corneal epithelium. (**D**) Limbal epithelial removal (white arrowheads) was performed after the formation of tdTomato^+^ stripe. Intravital tracking of accurate stripe was followed by fluorescent microscopy over time. (**E**) The large wound of central corneal epithelium was performed at 9 weeks after limbal epithelial removal, and the change of lineage-tracing tdTomato^+^ clone was investigated during the wound healing. (**F**) The whole-mount CK12 immunostaining of lineage-tracing corneas at 8 days after the large wound of central corneal epithelium. The limbus is localized based on the location of iris and ciliary body in the bright field of the whole-mount cornea. The “foot” means the end of corneal epithelial stripe that is closer to the limbus. Dashed circles show these clones. The arrows (**C**, **F**) point to CK12-negative de-differentiated LESCs. EL, eyelid; L, limbus; PC, peripheral cornea; CC, central cornea.

To investigate whether these tdTomato-labeled small clones de-differentiate into qLESCs, we performed wound healing with different-sized wounds as different responses of qLESCs to different-sized central corneal wounds (10, 30). After a small central corneal wound, the tdTomato-labeled clone neither proliferated nor migrated from the limbus, and was entirely CK12-negative (Fig. 4B, C), indicating the de-differentiation of CECs into qLESCs. After a medium central corneal wound, the tdTomato-labeled clone proliferated marginally and remained at the limbus. About half of the small clone was CK12^+^ CECs, while the other half was still CK12-negative (Fig. S4B, C). After a large central corneal wound, the tdTomato-labeled clone proliferated massively and migrated centripetally toward the central cornea. The “foot” of newly-formed stripe stayed at the limbus and was CK12-negative, while tdTomato-labeled cells that proliferated and migrated toward the central cornea were CK12^+^ CECs (Fig. 4E, F). Taken together, these results suggested the de-differentiation of CECs into qLESCs after the ablation of innate stem cells, and these qLESCs maintained quiescence during homeostasis and proliferated and differentiated into CECs to participate in wound healing after extensive wound.

### 3.4 No Trans-differentiation from Conjunctival Epithelial Cells

Recently, conjunctival epithelial cells were reported to be able to change into CK12^+^ cells with a stripe-like pattern in a total LSCD (tLSCD) model by using CK12 immunostaining (31). Thus, we examined whether conjunctival epithelial cells also contributed to the regeneration of LESCs after the ablation of innate LESCs through lineage tracing experiments. We found that some tdTomato-labeled clones and stripes from conjunctival epithelium emerged at the limbus after the ablation of innate LESCs in CK14-CreERT2;H11-reporter mice (Fig. 3A-C①③, Fig. 3D-F②; Fig. S5A). These clones and stripes didn’t invade the cornea during homeostasis; however, after a medium central corneal wound, they proliferated and migrated rapidly toward the central cornea, forming stripes. However, these tdTomato-labeled stripes consisted of CK12-negative cells (Fig. 3C①③, Fig. 3F②; Fig. S5A). Furthermore, we also performed a two-step removal of limbal and corneal epithelium to model tLSCD and didn’t found any CK12^+^ cells on the cornea after 1 month (Fig. S5B). Thus, these results ruled out the possibility that conjunctival epithelial cells trans-differentiated into LESCs or CECs to maintain the corneal epithelium.

### 3.5 The Outcome of the Competition between Conjunctival and Corneal Epithelium for the Limbus Determined the Occurrence of De-differentiation or Not

Confusingly, Park et al reported that cornea developed pathological features of LSCD, including corneal opacification, angiogenesis and the appearance of goblet cells within the injured zone, after a similar limbal epithelial removal (21). This implied that the de-differentiation of CECs might not occur under their experimental conditions. Thus, we next examined the process of the de-differentiation in detail to explore the potential reason for this apparent contradiction. At 12 hours after the ablation of innate LESCs, the tdTomato-labeled stripe moved obviously toward the limbus (Fig. 3A, D; Fig. 4A, D). This could be accomplished by one of three ways: 1) migration, 2) alteration in size and shape, or 3) proliferation of CECs. However, the F-actin staining and quantification of cellular area indicated that there was no obvious changes of cell shape and size of surrounding CECs after the removal of limbal epithelium (Fig. 5A, C), and the cell proliferation (EdU staining) was inhibited across the whole cornea at 12 hours when compared with normal homeostasis (Fig. 5B, D; Fig. S6). Therefore, it was highly likely that CECs migrated toward the limbus at the early stage as a response to the limbal epithelial removal. Subsequently, CECs proliferated rapidly to heal the limbal wound and even extended to conjunctival margin at 24 hours (Fig. 5B, D; Fig. S6; Fig. 2B). However, conjunctival epithelial cells proliferated markedly at 2 days. Thereafter, at 6 days, the proliferation of both corneal and conjunctival epithelial cells returned to their levels during normal homeostasis (Fig. 5B, D; Fig. S6). These results indicated that CECs rapidly responded to the limbal wound and covered the limbus earlier than conjunctival epithelial cells when the de-differentiation of CECs could occur.

**Figure 5:**
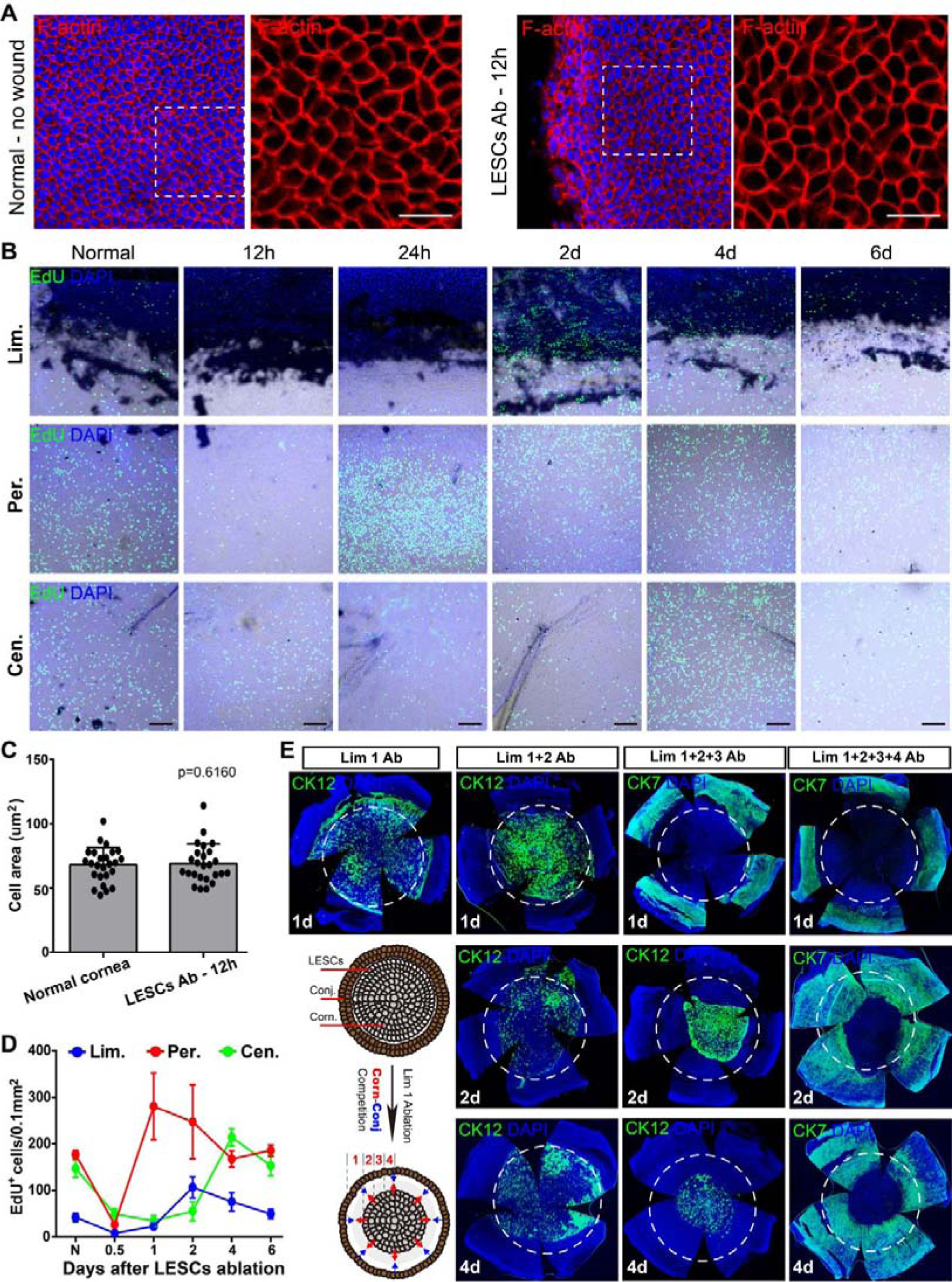
The competition between conjunctival and corneal epithelium for the limbus determined the occurrence of de-differentiation or not. (**A**, **C**) The whole-mount F-actin staining of normal cornea and cornea at 12h after the removal of limbal epithelium. The cellular area of peripheral corneal epithelial cells was quantified. Data are the mean ± SD, n=25 cells; statistical analysis are performed by Student’s unpaired t-test, and p value is shown. (**B**, **D**) The EdU-positive proliferative epithelial cells at limbus (Lim.), peripheral cornea (Per.) and central cornea (Cen.) at indicated time points after the removal of limbal epithelium were quantified. Data are the mean ± SD, n=8 zones. (**E**) The ocular surface between the conjunctiva and central cornea was regionalized into four parts, namely 1, 2, 3, and 4 as shown. The different-sized zones of limbal and corneal epithelium were scratched off, resulting in different outcomes of the competition between corneal and conjunctival epithelial cells for the limbus. The whole-mount CK12 or CK7 immunostaining was performed against corneas with different-sized wounds. Dashed circles indicate the limbus. Scale bars, 20um (**A**) and 100um (**B**).

As known, limbal niche plays key role in governing the fate of LESCs (17). We found that the removal of limbal epithelium in Park’s study extended to the peripheral cornea, which was a larger wound when compared with ours. This implies that CECs may not occupy the limbus and de-differentiate into LESCs under Park’s experimental conditions, because conjunctival cells will migrate to and occupy the limbus earlier than CECs. Therefore, we hypothesized that the competition between corneal and conjunctival epithelium for the limbus would determine which cells occupy the limbus and the de-differentiation of CECs occurred or not. To test this hypothesis, we wounded the limbal and corneal epithelium with varying sizes, resulting in either corneal or conjunctival epithelial cells occupy the limbus (Fig. 5E). In the (Lim 1) ablation, CECs covered the limbus at 24 hours, de-differentiated into LESCs and maintained the function of corneal epithelium as shown in our above data. In the (Lim 1+2) ablation, most zones of limbus were covered by CECs and the cornea maintained normal function. However, in the (Lim 1+2+3) and (Lim 1+2+3+4) ablations, CECs never covered the limbus, leading to the corneal conjunctivalization (Fig. 5E). Thus, these results suggested that corneal and conjunctival epithelial cells competed for the limbal niche, and the occupy of the limbus by CECs was the prerequisite for the occurrence of de-differentiation.

### 3.6 The De-differentiation of CECs Required an Intact Limbal Niche

Since the cover of CECs onto the limbus determined the de-differentiation of CECs would occur or not, we then explored whether an intact limbal niche was required for the de-differentiation. We scratched off the whole limbal epithelium to ablate innate LESCs, and then applied NaOH (1M, 5s) to a localized zone of the limbus (approximately a quarter of the whole limbus) that would cause the saponification of fatty acids in cell membranes and the destruction of proteoglycan ground substance and destroyed the limbal niche (32). This experimental strategy allowed us to be able to compare the fate of CECs with or without an intact limbal niche on the same cornea (Fig. 6A).

**Figure 6:**
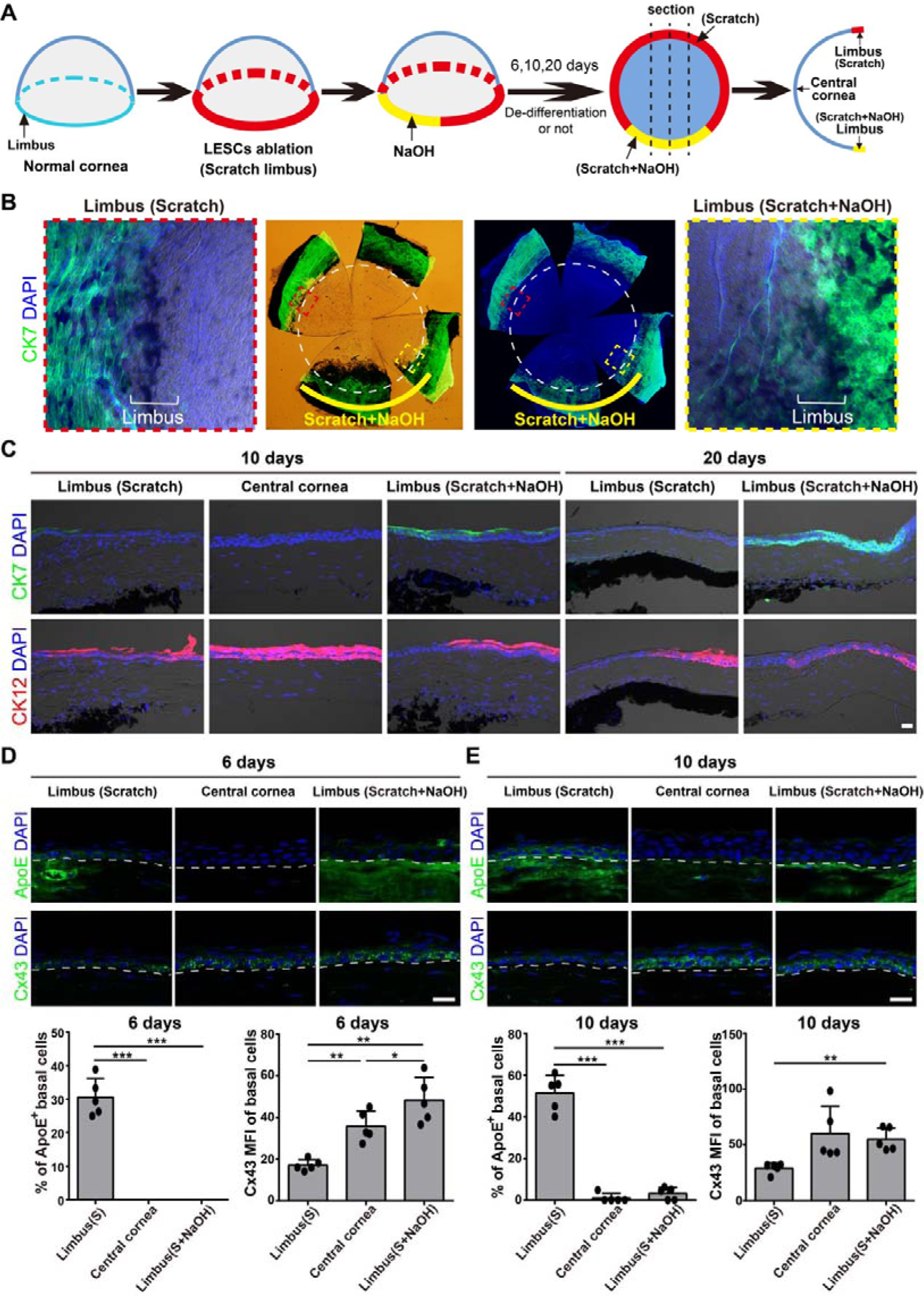
The de-differentiation of CECs required an intact limbal niche. (**A**) Experimental strategy of the localized NaOH application to destroy the limbal niche, which allows the examination of the fate of CECs either with or without an intact limbal niche on the same cornea. (**B**) The whole-mount CK7 immunostaining against corneas of scratched limbus either with or without NaOH application. Dashed circles indicate the limbus. (**C**) The frozen-section CK7 and CK12 immunostaining against cornea of scratched limbus either with or without NaOH application at 10 and 20 days. (**D, E**) The frozen-section Cx43 and ApoE immunostaining against cornea of scratched limbus either with or without NaOH application at 6 and 10 days. The percentage of ApoE^+^ LESCs and the mean fluorescence intensity (MFI) of Cx43 were quantified. Data are the mean ± SD, n=5 biological replicates; statistical analysis were performed by paired one-way ANOVA with Tukey’s test. Scale bars, 20 um.

At 20 days after the injury, conjunctival epithelial cells invaded the cornea across the limbus where both scratch and NaOH were applied, but not through the limbus where only a scratch was applied (Fig. 6B; Fig. S7A). At the limbus where only scratch was applied, basal CECs reduced the level of CK12 at 10 days and lost its expression at 20 days (Fig. 6C). Conversely, at the limbus where both scratch and NaOH were applied, CECs maintained a high CK12 level at 10 days, and were replaced by conjunctival epithelial cells at 20 days (Fig. 6C). Consistent with previous results, at the limbus where only scratch was applied, CECs expressed ApoE and showed low Cx43 expression at 6 and 10 days (Fig. 6D, E), indicating the de-differentiation of CECs. However, at the limbus where both scratch and NaOH were applied, CECs didn’t express ApoE and maintained the high Cx43 level at 6 and 10 days (Fig. 6D, E), indicating the persistence of CECs’ fate and the absence of their de-differentiation into LESCs. Taken together, all these results suggested that the destruction of the limbal niche by the NaOH application impeded the de-differentiation of CECs, and that the de-differentiation required an intact limbal niche.

### 3.7 The Hippo/YAP Pathway Mediated the De-differentiation of CECs

To investigate the potential role of Hippo/YAP pathway in the de-differentiation of CECs into LESCs, we first examined the expression and intracellular distribution of YAP in corneal epithelium of mice. During homeostasis, a higher expression level of YAP was found in peripheral and central corneal epithelium compared to the limbal epithelium. YAP in most limbal epithelial basal cells were cytoplasmic, and rare limbal epithelial basal cells were evenly nuclear and cytoplasmic (N/C) or nuclear YAP. However, most basal cells of peripheral and central epithelial cells were N/C and even nuclear YAP (Fig. S8A). This indicated the higher YAP activity in peripheral and central epithelial cells than in limbal epithelial cells, which was further confirmed by expressions of inactive YAP (phosphorylated YAP at Ser127) and active YAP (non-phosphorylated YAP) (Fig. S8B, C). This observation is consistent with the pro-proliferative activity of YAP and the higher proliferative rate of peripheral and central epithelial cells than limbal epithelial cells (Fig. 5B, D, Fig. S6).

After the ablation of limbal epithelium, YAP in CECs at the limbus rapidly translocated into the nucleus (active YAP) from 12 hours to 2 days. Subsequently, the expression level and activity of YAP decreased and returned to homeostatic levels at 6 days when LESCs regenerated (Fig. 7A; Fig. S9A, B). The dynamic of YAP activity was in line with the proliferation of CECs during de-differentiation (Fig. 5B, D, Fig. S6). These results indicated that YAP was re-activated with increased proliferation of CECs at the early stage and recovered to a low level and activity as CECs de-differentiated. Then, we administered verteporfin (VTP, an inhibitor of YAP-TEAD activity) or TRULI (an inhibitor of LATS, indirectly activating YAP)(33) onto the cornea for 4 days and examined the de-differentiation of CECs at 6 days. The administration of VTP and TRULI respectively decreased and increased YAP activity and cell proliferation at the limbus (Fig. S9C). VTP significantly increased the ratio of ApoE^+^ LESCs and reduced the expression of Cx43 in limbal basal cells, indicating the enhanced de-differentiation of CECs. In contrast, TRULI inhibited the de-differentiation of CECs (Fig. 7B). Together, these results suggested that the LATS-dependent suppression of YAP promoted the de-differentiation of CECs into LESCs *in vivo*.

**Figure 7:**
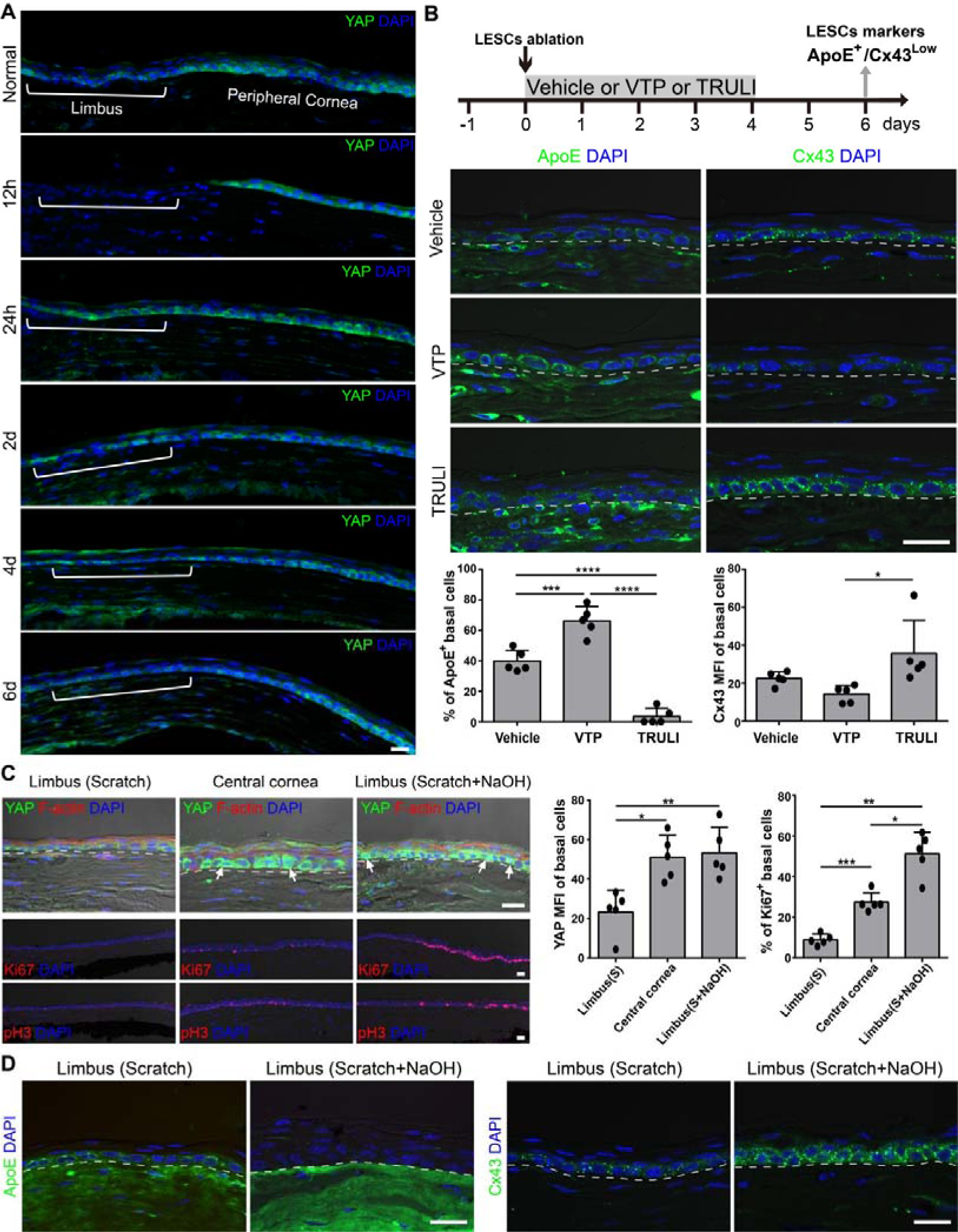
The Hippo/YAP pathway mediated the de-differentiation of CECs. (**A**) Immunostaining of YAP in frozen sections of normal cornea and LESCs-ablation cornea at indicated days after the limbal epithelium removal. The limbus was shown. (**B**) The administration of vehicle or VTP or TRULI on cornea for 4 days after the LESCs ablation, and expressions of ApoE and Cx43 were examined by immunostaining at 6 days. The percentage of ApoE^+^ LESCs and the mean fluorescence intensity (MFI) of Cx43 in limbal basal cells were quantified. (**C**) The frozen-section YAP, Ki67 and pH3 (indicators of cell proliferation) immunostaining against cornea of scratched limbus either with or without NaOH application at 10 days after the LESCs ablation. Arrows point to CECs with nuclear YAP. The MFI of YAP in limbal basal cells and the percentage of Ki67^+^ limbal basal cells were quantified. (**D**) Corneas with scratched limbus and localized NaOH application were administrated with VTP for 4 days. The frozen-section ApoE and Cx43 immunostaining against cornea of scratched limbus either with or without NaOH application at 6 days after the LESCs ablation. Data are the mean ± SD, n=5 biological replicates; statistical analysis were performed by unpaired (**B**) or paired (**C**) one-way ANOVA with Tukey’s test. Scale bars, 20 um.

Next, we studied the YAP signal in CECs where both scratch and NaOH were applied to the limbus, which didn’t de-differentiate into LESCs (Fig. 6D, E). As expected, CECs at limbus where both scratch and NaOH were applied maintained high level and activity of YAP (Fig. 7C; Fig. S7B). Consistent with the high YAP level and activity, CECs at limbus where both scratch and NaOH were applied rapidly proliferated (Fig. 7C; Fig. S7B). In contrast, YAP level and activity of CECs at limbus where only scratch was applied decreased and recovered to that level during homeostasis when LESCs regenerated at limbus (Fig. 6D, E; Fig. 7C; Fig. S7B). Together with above results, these data indicated that the persistent high level and activity of YAP hindered the de-differentiation of CECs at limbus where both scratch and NaOH were applied. Then, we examined whether inhibiting YAP activity by VTP could rescue the capacity of CECs to de-differentiate into LESCs and prevent LSCD. Disappointingly, VTP administration didn’t restore the de-differentiation of CECs into LESCs (Fig. 7D), and CECs still maintained the high YAP level and proliferative rate (Fig. S7C). These results suggested that the high YAP level and activity of CECs after NaOH burn to the limbus hindered their de-differentiation, and inhibition of YAP-TEAD via VTP couldn’t rescue the capacity of CECs to de-differentiate into LESCs.

## 4. Discussion

In injured tissues, regeneration is often associated with cell fate plasticity (de-differentiation and trans-differentiation), where cells deviate from their normal lineage paths (34). It is becoming clear that this plasticity creates alternative strategies to restore damaged or lost cells and will initiate a new class of therapies by modulating cell fate (35). In this study, we showed that CECs de-differentiated into functional LESCs, which contributed to the functional maintenance of corneal epithelium, after innate stem cell ablation. This might open up an alternative path for the efficient treatment of LSCD by modulating the de-differentiation of CECs *in vivo*.

### 4.1 The de-differentiation of CECs into LESCs

A previous study showed that cornea-committed epithelial cells could travel in the opposite direction (i.e., centrifugally), convert into CK15-GFP^+^ LESCs and reestablish the tissue boundary (20). However, recent studies have challenged the use of CK15 as an ideal marker for LESCs and refuted the phenomenon of de-differentiation and even the centrifugal migration of CECs (12, 21). In recent years, an increasing number of researchers proposed to define adult stem cells by their biological functions rather than markers (26, 27). In corneal epithelium, LESCs are endowed with three vital functions: 1) maintaining corneal epithelial homeostasis, 2) forming a boundary barrier to avoid conjunctival epithelial invasion (i.e. corneal conjunctivalization), and 3) participating in wound healing by rapid proliferation and migration. Herein, we showed the maintenance of corneal epithelium, the lack of corneal conjunctivalization and the normal corneal epithelial wound healing after the ablation of innate stem cells, indicating the regeneration of functional LESCs. By employing lineage tracing, intravital imaging and immunostaining, we highlighted the contribution of the de-differentiation of CECs, but not trans-differentiation of conjunctival epithelial cells, to these regenerated LESCs. More importantly, we postulated the competition between corneal and conjunctival epithelium for the limbus, which determined which cells occupied the limbus and whether de-differentiation would occur or not. The de-differentiation only occurred when CECs outcompeted conjunctival epithelial cells and occupied the limbus. Thus, the sizes of the ring-shaped limbal wounds will determine the outcomes of the competition between corneal and conjunctival epithelium for the limbus, determining whether de-differentiation occurs and ultimately the final fate of the cornea. Thus, our results largely reconciled the apparent contradictions in the field of corneal epithelial de-differentiation.

Although our results suggest the capacity of CECs to de-differentiate into LESCs *in vivo*, there are still many questions to be solved. First, it is unclear whether all committed or differentiated CECs can undergo successful de-differentiation into LESCs, or whether this is limited to a small sub-population (e.g., corneal basal TACs). In the airway epithelium, the ability of differentiated cells to acquire stem cell properties is inversely proportional to the degree of the maturity of differentiated cells (36). As known, the hierarchy also exists in the CECs, e.g. the proliferative capacity (5). Further studies are needed to investigate whether the hierarchy of CECs also influences the propensity of de-differentiation. Second, although we demonstrate the contribution of de-differentiation of CECs to the regeneration of LESCs and rule out the conjunctival epithelial trans-differentiation, other origins may also exist. For instance, limbal mesenchymal stem cells have been reported to express not only mesenchymal stem cells markers but also ABCG2, ABCB5, ALDH3A1, PAX6 and p63α that were specific for LESCs, and differentiate into epithelial cells (37, 38). Third, the de-differentiation of CECs requires the limbal niche. A recent study showed that CD90^+^/CD105^+^/SCF^+^/PDGFRβ^+^ limbal niche cells promoted the de-differentiation of CECs in three-dimensional Matrigel (39). How do limbal niche cells regulate the de-differentiation of CECs? Whether transplantation of limbal niche cells could prevent LSCD after NaOH burn via rescuing the de-differentiation of CECs? Whether other niche factors, such as immune cells and extracellular matrix (ECM), also play roles in the de-differentiation of CECs? All these questions need to be solved in the future. Fourth, limbal removal could cause mild neovascularization in 4 out of 12 rabbits and subsequently corneal injury induced a higher frequency and severity of LSCD (40). Thus, the de-differentiation in cornea of rabbit, and potentially humans, is still elusive and requires further study. Addressing these questions will pave the way to develop potential therapies for related human corneal diseases.

It should be noted that the de-differentiation of CECs into aLESCs or qLESCs occurred in lineage-tracing mice driven by different promoters. Actually, almost all de-differentiation of CECs into aLESCs occurred in the CK14-induced lineage-tracing mice, while qLESCs in Slc1a3-induced lineage-tracing mice. This implies that the interference of the expression of endogenous CK14 and Slc1a3 might affect the process of de-differentiation and the final fate of CECs. The use of non-specific-promoter-driven lineage-tracing technique will resolve this limitation of our study.

### 4.2 The trans-differentiation from conjunctival epithelial cells

In the 1980s, the trans-differentiation of conjunctival epithelial cells into CECs was solidified, where a large chemical (n-heptanol) injury was applied to the corneal epithelium (which likely extended to the limbus through lateral diffusion) and conjunctival epithelial cells changed morphologically into cornea-like epithelia to maintain transparency of cornea (41, 42). Then, the trans-differentiation of conjunctival epithelial cells was found in more experimental conditions (43–45). However, the notion of conjunctival trans-differentiation into CECs was revoked later, as the possibility of incomplete removal of the limbal epithelium and/or its LESCs population (46). Recently, the trans-differentiation of conjunctival epithelial cells into CECs was re-solidified, where the limbal and corneal epithelium were totally removed, and conjunctival cells ventured over the cornea and acquired the capacity to trans-differentiate into K12^+^ epithelia (31, 47).

In this study, to examine whether conjunctival epithelial cells also contribute to the regeneration of LESCs after the ablation of innate LESCs, we traced tdTomato-labeled clones from conjunctival epithelium in CK14-CreERT2;H11-reporter mice. We found that these conjunctival clones and stripes didn’t invade the cornea during homeostasis, and they ventured over the cornea after a central corneal wound but are still CK12-negative cells. Furthermore, we also performed a two-step removal of limbal and corneal epithelium to model tLSCD and didn’t found any CK12^+^ cells on the cornea after 1 month. Thus, our results ruled out the possibility that conjunctival epithelial cells trans-differentiated into LESCs or CECs to maintain the corneal epithelium. One fact should be noted is that these evidences that supported the notion of conjunctival trans-differentiation were not based on lineage tracing. Thus, they cannot rule out the possibility of residual LESCs and/or CECs contribute to these K12^+^ epithelia. Of course, the CK14-induced lineage-tracing mice might also affect the process of trans-differentiation of conjunctival epithelial cells. Thus, the non-specific-promoter-driven lineage-tracing technique should be used to further explore the possibility of conjunctival trans-differentiation. In addition, other cells, but not conjunctival and corneal epithelial cells, should also be considered to contribute to these K12^+^ epithelia.

### 4.3 The role of YAP in the de-differentiation of CECs

During the submission of our manuscript, Bhattacharya et al. reported that the activity of YAP played significant roles in maintaining LESCs via the mechanotransduction pathway, and the inhibition of YAP by VTP hindered the de-differentiation of CECs *in vivo* (48). However, we observed rare nuclear YAP in LESCs during normal homeostasis. Instead, we detected higher YAP level and activity in peripheral and central epithelial cells, but not in LESCs. Confusingly, previous studies reported apparent contradictions in YAP location and activity, where higher activity of YAP was found either at the limbus (24) or in peripheral and central cornea (49). We believe that the specific epitope of the antigen to which the antibody binds may influence these results. Some YAP antibodies bind to the epitope of YAP including S127 or S381 site, which are crucial for the regulation of YAP location and activity. Some YAP antibodies may also bind to paralog WWTR1/TAZ, which has similar sequence of amino acid and function with YAP. In addition, the type of experimental animal should also be considered, as the structures of the limbus in different animals are different. As for the opposing effects of VTP on the de-differentiation of CECs, we surmise that might result from these different markers used to identify LESCs. Bhattacharya et al. used CK15-GFP, CD63 and IFITM3 to mark LESCs (48). However, we found that CD63 and IFITM3 were also expressed by basal cells of peripheral and central epithelium. Instead, we tested various markers and ultimately selected ApoE^+^/Cx43^low^/CK12^-^ to define LESCs. Interestingly, CK15-GFP was a useful indicator of aLESCs in previous reports (11, 48), while ApoE was mainly expressed in qLESCs as shown in single cell RNA sequencing (48). Thus, we wonder whether the opposing effects of VTP on the de-differentiation of CECs may reflect the identification of different sub-populations of LESCs (aLESCs and qLESCs). If so, the administration of VTP might inhibit the regeneration of aLESCs, and promote the conversion of aLESCs into qLESCs, which aligns with the essential function of YAP in promoting cell proliferation. This hypothesis was indirectly supported by a previous study, where the lineage tracing was induced by CK12 promoter and deletion of YAP reduced stripes formed by aLESCs via the inhibition of CECs’ proliferation (24). Moreover, it should be noted that persistent YAP activation appears to be unfavorable for the de-differentiation of CECs and may even cause their trans-differentiation into epidermal epithelium under chronic inflammation (50, 51). Therefore, the roles of YAP in the maintenance of LESCs, the conversion between LESCs sub-populations, the de-differentiation of CECs and the trans-differentiation of corneal epithelium needs further investigations.

The cell plasticity or fate change is coordinated by both intrinsic changes within the cell and extrinsic microenvironment. Here, we reported that the LATS-dependent suppression of YAP promoted the de-differentiation of CECs into LESCs after the scratch off of limbal epithelium, while persistent YAP activation of CECs at the limbus after NaOH treatment inhibited the de-differentiation of CECs. The activity of the Hippo pathway is regulated by a network of upstream inputs involved in cell adhesion, cell junction, cell morphology and cell polarity. In addition, the Hippo pathway is also modulated by crosstalk with other signaling pathways, such as Wnt, and YAP can interact with other transcription factors such as p73 and RUNX. This might explain why VTP (an inhibitor of YAP-TEAD interaction) couldn’t rescue the de-differentiation of CECs after NaOH burn, as in the case of NaOH burn, YAP might interact with other transcription factors but not TEAD. One factor that should be noted is the stiffness of limbal stroma. The matrix of central cornea was much stiffer than the limbus, and the activity of YAP was shown to change when cells grew upon substrates of increasing stiffness (52). Studies have shown that NaOH burn increased the stiffness of cornea (32, 50), which might hinder the de-differentiation of CECs via YAP-mediated mechanotransduction. Importantly, stiffer stroma increases YAP activity via FAK and SRC kinases or actin cytoskeleton independently of LATS (53–56). Whether the increasing stiffness of limbal stroma induce the high YAP activity of CECs after NaOH burn? How did the high YAP activity inhibit the de-differentiation of CECs into LESCs? The better understanding of their underlying molecular mechanisms are much-needed and would promote the development of effective treatments for LSCD by modulating the fate of CECs.

### 4.4 Limitations of this study

First, this study shows the de-differentiation of CECs, but not trans-differentiation of conjunctival cells, into LESCs. However, it don’t completely rule out the possibility that residual LESCs contribute to the regeneration of LESCs, even if it shows that there are no residual epithelial cells at the limbus. Second, the de-differentiation of CECs into aLESCs or qLESCs occurs in lineage-tracing mice driven by different promoters. In the future, the non-specific-promoter-driven lineage-tracing technique will be used to resolve this limitation. Third, only the co-factor YAP of Hippo pathway is examined in the study, and the role of its paralog TAZ should be explored in further. In addition, inhibitors of YAP-TEAD and LATS kinase are used in this study, and gene knockout should be considered as a powerful method to explore the role of Hippo/YAP pathway in the de-differentiation in the future.

## Grant & funding information

This work is supported by grants from National Natural Science Foundation of China (81900832, 81970843), National Key R&D Program of China (2018YFA0107302), and the Science and Technology Innovation Project of Army Medical University (2020XQN14).

## Author contributions

Y. Li, Y. Liu, and H. Xu designed the project. Y. Li wrote the manuscript and performed most of these experiments, including the LESCs ablation model, corneal epithelial wound healing and drug treatment, lineage tracing and intravital imaging, and fluorescence staining. L. Ge performed fluorescence staining, B. Ren and Z. Yin performed the cross of transgenic mice, X. Zhang and Y. Yang performed the intravital imaging, and H. Liu made important suggestions on the design of some experiments and the first draft of the manuscript. All authors read and approved the final manuscript.

## Declaration of competing interest

The authors declare no competing interests.

## Data availability

Data will be made available on request.

## Notes

### Competing Interest Statement

The authors have declared no competing interest.

