## Supplementary figures for "The Hippo/YAP Pathway Mediates the De-differentiation of Corneal Epithelial Cells into Functional Limbal Epithelial Stem Cells *In Vivo*"

**Figure S1: The recovery of the function of limbus after innate LESCs ablation *in vivo*.**

**Figure S2: The recovery of the function of limbus after innate LESCs ablation in rat model.**

**Figure S3: The examination of lineage tracing mice (CK14-CreERT; H11-reporter).**

**Figure S4: The response of de-differented qLESCs to medium wound of central corneal epithelium.**

**Figure S5: Regenerated LESCs were not from trans-differentiation of conjunctival epithelial cells.**

**Figure S6: The cell proliferation of corneal epithelium after the ablation of innate LESCs.**

**Figure S7: The de-differentiation of CECs required an intact limbal niche and the suppression of YAP signal.**

**Figure S8: The expression of YAP in cornea during normal homeostasis.**

**Figure S9: The expression of YAP at limbus after the ablation of innate LESCs.**

**Figure S10: The model of the de-differentiation of CECs.**

### Supplementary figures and legends

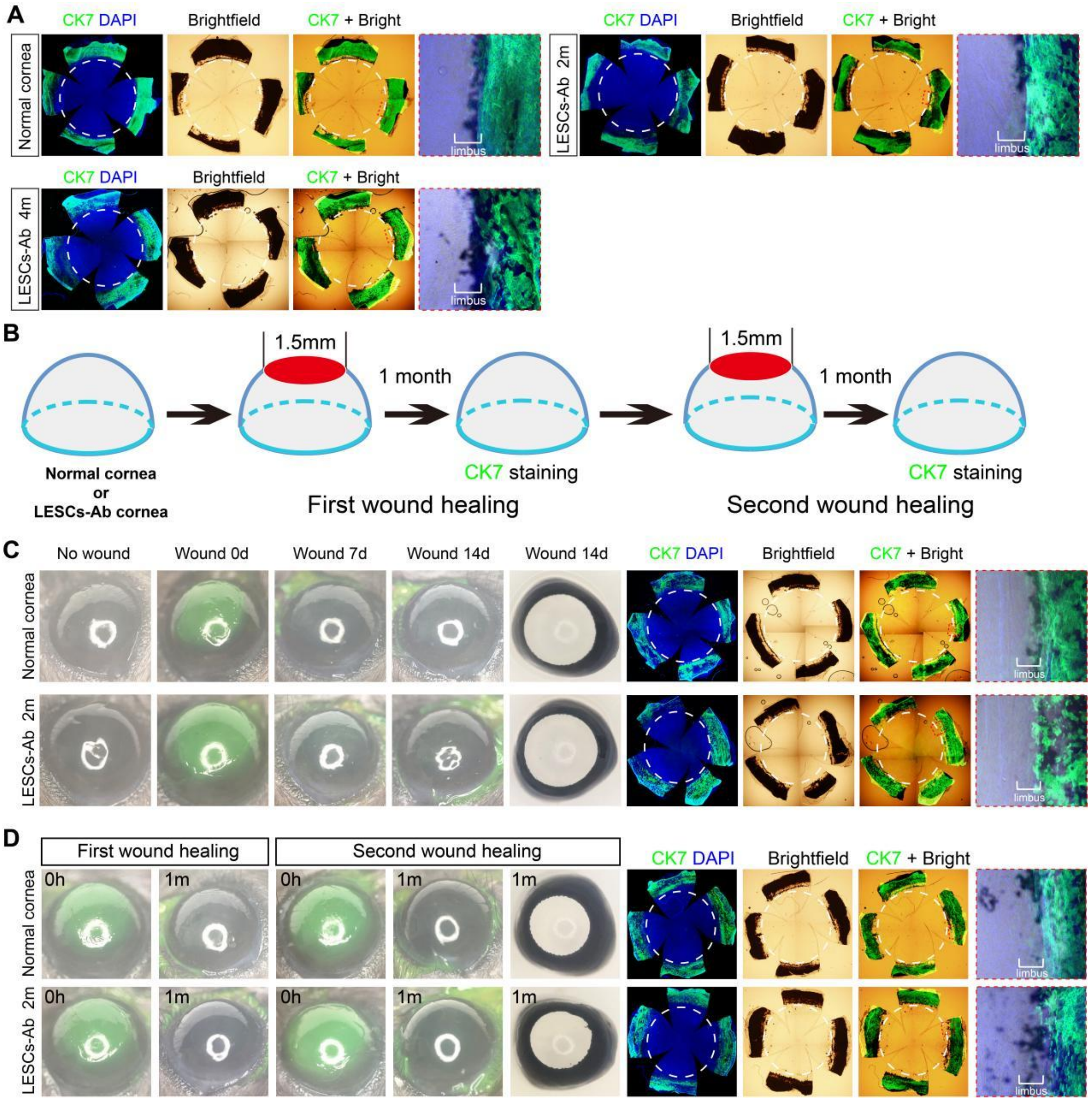


**Figure S1: The recovery of the function of limbus after innate LESCs ablation *in vivo*.**

(**A**) The limbal epithelium and marginal corneal and conjunctival epithelium of 6-month-old C56BL/6J mice were removed by surgery. Whole-mount immunostaining of normal and LESCs-ablation corneas for conjunctival epithelial marker CK7 at 2 and 4 months after limbal epithelial removal. (**B**) Experimental strategy of two consecutive central corneal wound healing of normal and LESCs-ablation corneas. CK7 staining was performed to examine whether conjunctival epithelial cells invaded the cornea. (**C**, **D**) The first and second wound healing of normal and LESCs-ablation corneas. Fluorescein sodium images and whole-mount staining of normal and LESCs-ablation corneas for CK7 are shown. Dashed circles indicate the limbus, which is localized based on the location of iris and ciliary body in the bright field.These results suggest that there is no corneal epithelial conjunctivalization on LESCs-ablation cornea even after two consecutive central corneal wound healing.

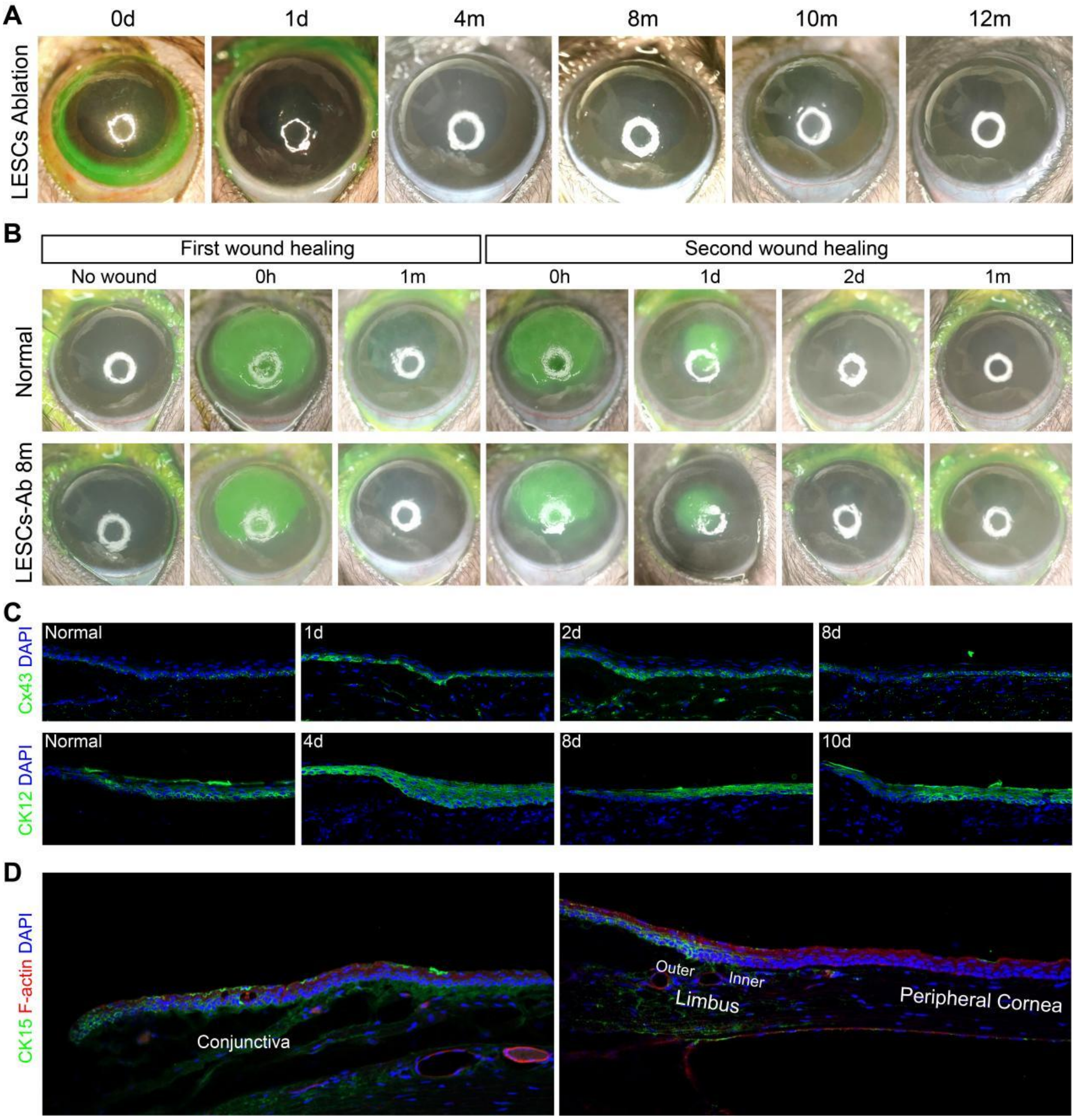


**Figure S2: The recovery of the function of limbus after innate LESCs ablation in rat model**.

(**A**) The limbal epithelium and marginal corneal and conjunctival epithelium of 6-month-old Long Evans rats were removed by surgery. Fluorescein sodium staining and bright-field images of injured eyes at indicated days. The corneas maintained transparency for at least 12 months. (**B**) Two consecutive central corneal wound healing of normal and LESCs-ablation corneas of rats was performed, and fluorescein sodium staining of injured eyes was imaged at indicated days. LESCs-ablation corneas showed identical healing rates to that of normal corneas during both first and second central corneal wound healing. (**C**) Immunostaining of Cx43 and CK12 in normal cornea and LESCs-ablation corneas at indicated days after limbal epithelium removal. Epithelial cells at the limbus showed reduced Cx43 and CK12 at 8 days, indicating the regeneration of LESCs after the ablation of innate LESCs. (**D**) Immunostaining of CK15 in normal cornea and conjunctiva of rat. CK15 was expressed in outer LESCs and conjunctival epithelium, and the antibody we used couldn't detect CK15 in cornea of mice. Thus, CK15 was not used as a marker of LESCs in this study.

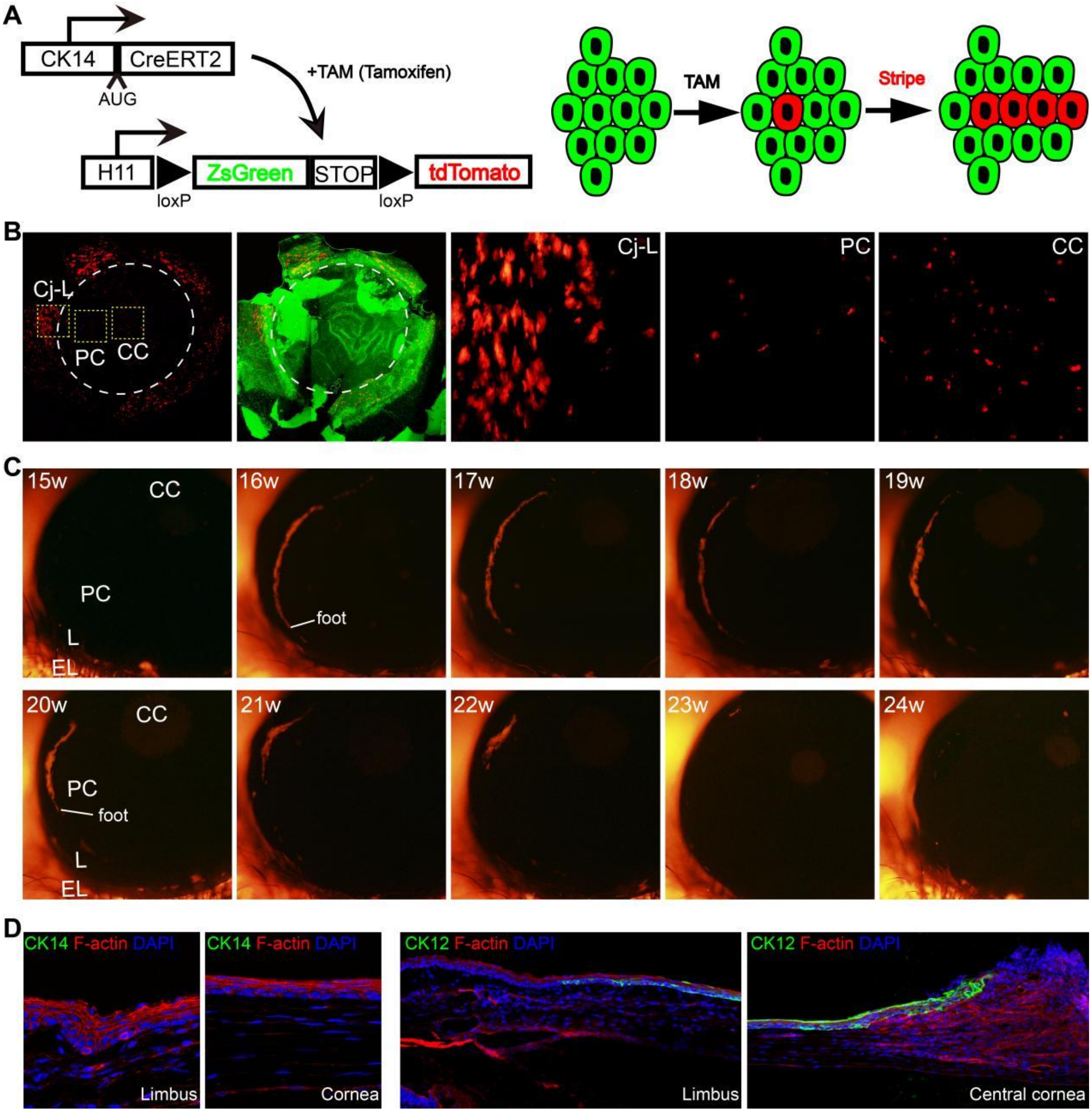


**Figure S3: The examination of lineage tracing mice (CK14-CreERT; H11-reporter).**

(**A**) Lineage tracing was induced by tamoxifen in 6-month-old mice (CK14/Slc1a3-CreERT; H11-reporter). These tdTomato-labeled LESCs or TACs proliferate and migrate centripetally toward the central cornea, forming stripes. Note that CreERT2 was knocked in the endogenous CK14 gene, while CreERT was under the control of exogenous Slc1a3 promoter that randomly integrated into the genome. (**B**) The tdTomato-labeled cells (CK14-CreERT2; H11-reporter) covered the whole ocular surface after tamoxifen induction for 7 days. Dashed circles indicate the limbus. Note that conjunctival epithelium was also labeled. (**C**) A typical corneal epithelial stripe that resulted from aLESCs after tamoxifen induction. Because of the rule of stochastic competition and neutral drift, this tdTomato-labeled corneal epithelial stripe gradually disappeared as the neutral competition between labeled LESCs and non-labeled LESCs. The "foot" means the end of corneal epithelial stripe that is closer to the limbus. EL, eyelid; Cj-L, conjunctiva-limbus; L, limbus; PC, peripheral cornea; CC, central cornea. (**D**) Immunostaining of CK14 and CK12 in corneal sections of CK14-CreERT2 homozygous mice. As the destruction of endogenous CK14 gene, the CK14-CreERT2 homozygous mice were similar to that of CK14 gene knock-out mice. CK14 immunostaining indicated the lack of the expression of CK14. In addition, the limbus and central cornea of CK14-CreERT2 homozygous mice showed to be abnormal.

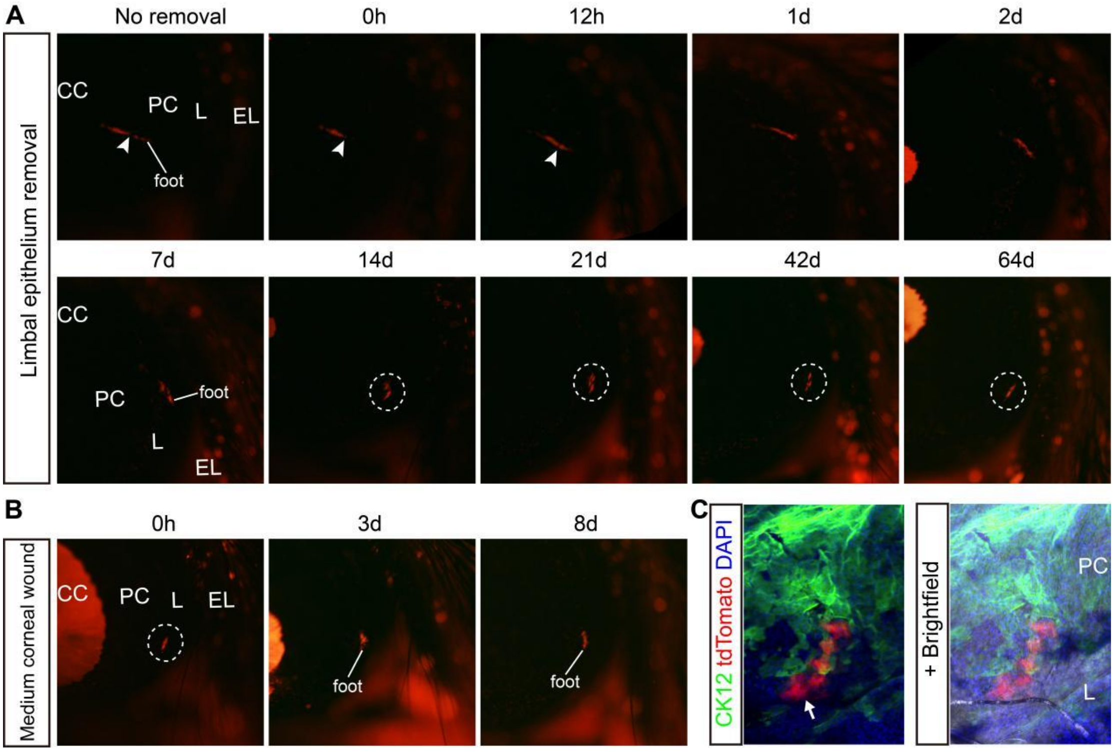


**Figure S4: The response of de-differented qLESCs to medium wound of central corneal epithelium.**

(**A**) The 6-month-old transgenic mouse (Slc1a3-CreERT; H11-reporter) was induced to express tdTomato reporter and limbal epithelial removal (white arrowheads) was performed after the formation of tdTomato$^{+}$ stripe. Intravital tracking of accurate stripe was followed by fluorescent microscopy over time. (**B**) The central corneal epithelium was scratched off (medium wound) at 64 days after limbal epithelial removal, and the change of lineage-tracing tdTomato$^{+}$ clone was investigated during the wound healing. (**C**) The whole-mount CK12 immunostaining of lineage-tracing cornea at 8 days after the medium wound of central corneal epithelium. The "foot" means the end of corneal epithelial stripe that is closer to the limbus. Dashed circles show these clones. The arrow points to CK12-negative de-differentiated LESCs. EL, eyelid; L, limbus; PC, peripheral cornea; CC, central cornea.

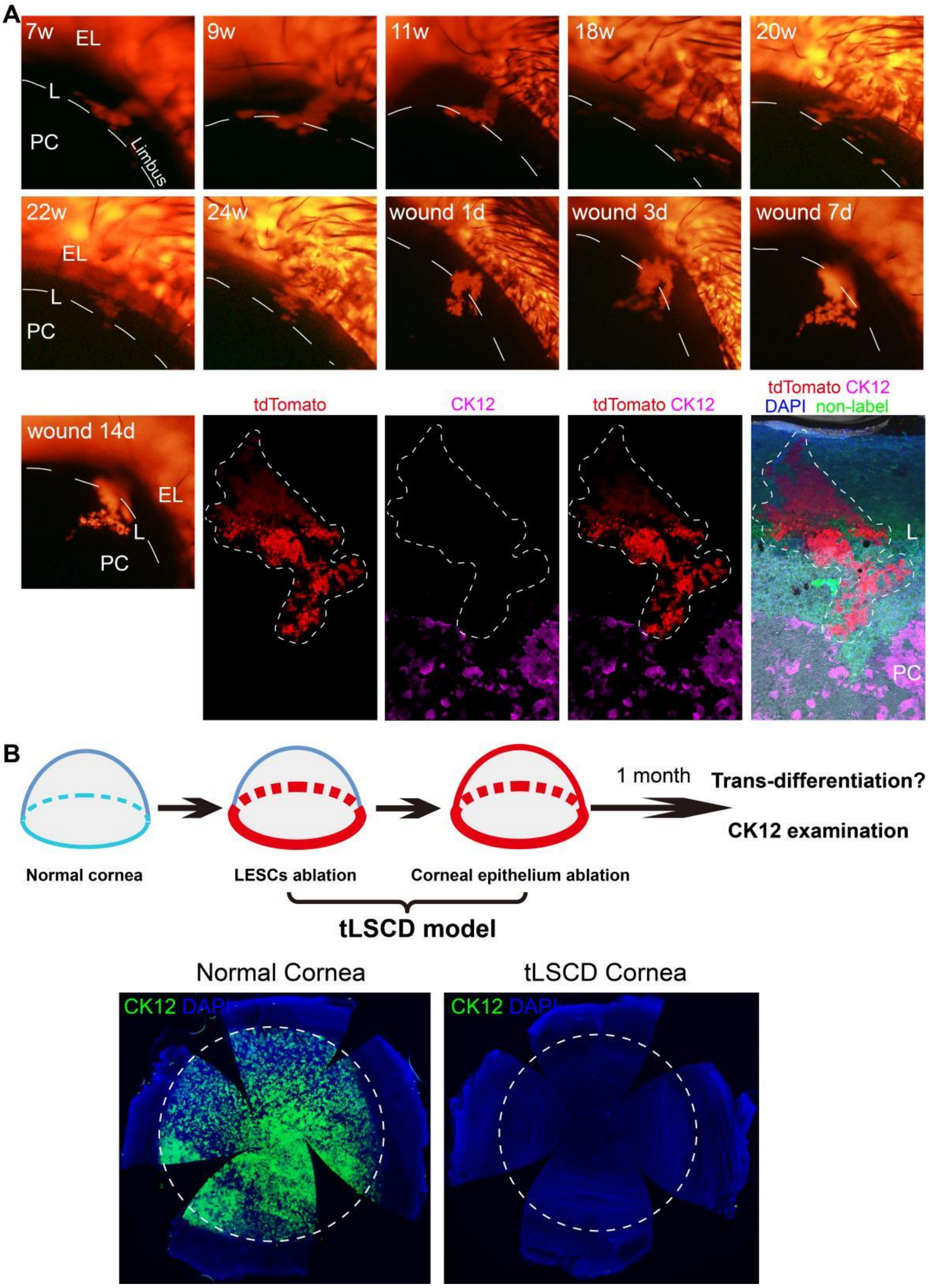


**Figure S5: Regenerated LESCs were not from trans-differentiation of conjunctival epithelial cells.**

(**A**) The dynamic changes of a conjunctival epithelial stripe after limbal epithelial removal for 24 weeks and central corneal epithelial wound healing for 14 days. Dashed lines indicate the limbus. The result of whole-mount CK12 immunostaining ruled out the possibility that conjunctival epithelial cells trans-differentiated into LESCs or CECs. (**B**) Two-step removal of limbal and corneal epithelium to model total LSCD (tLSCD), and CK12 staining of normal and tLSCD corneas was performed after 1 month. The normal cornea was used as control. Dashed circles indicate the limbus. The absence of $CK12^{+}$ CECs indicated that conjunctival epithelial cells were not able to trans-differentiate into $CK12^{+}$ CECs to maintain the function of cornea. EL, eyelid; L, limbus; PC, peripheral cornea.

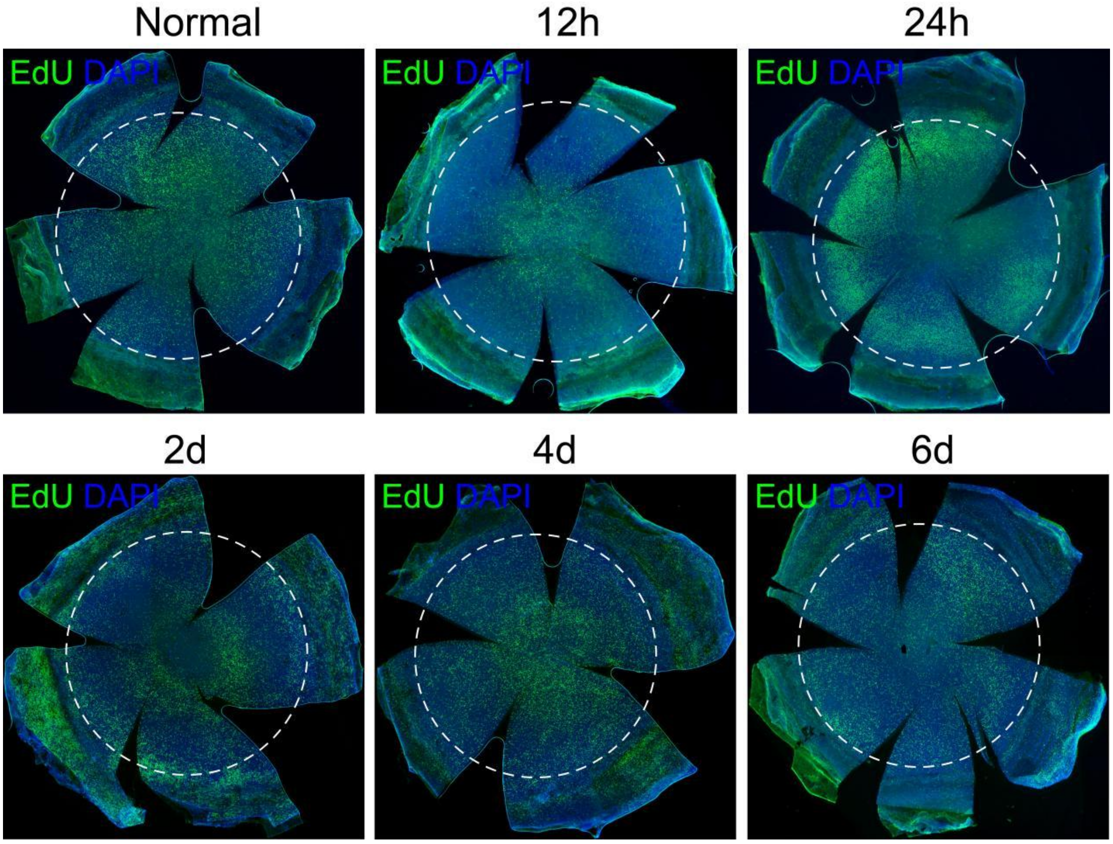


**Figure S6: The cell proliferation of corneal epithelium after the ablation of innate LESCs.**

The cell proliferation across the whole ocular surface was detected by EdU staining at indicated days after the ablation of innate LESCs. Note that the cell proliferation across the whole ocular surface was inhibited at 12h after the ablation of innate LESCs when compared with normal homeostasis. The increased cell proliferation of conjunctival epithelium occurred at 2 days after the ablation of innate LESCs when CECs have covered the limbus and even extended to conjunctiva. The proliferation of both corneal and conjunctival epithelial cells at 6 days after limbal epithelial removal approximately returned to their levels during normal homeostasis.

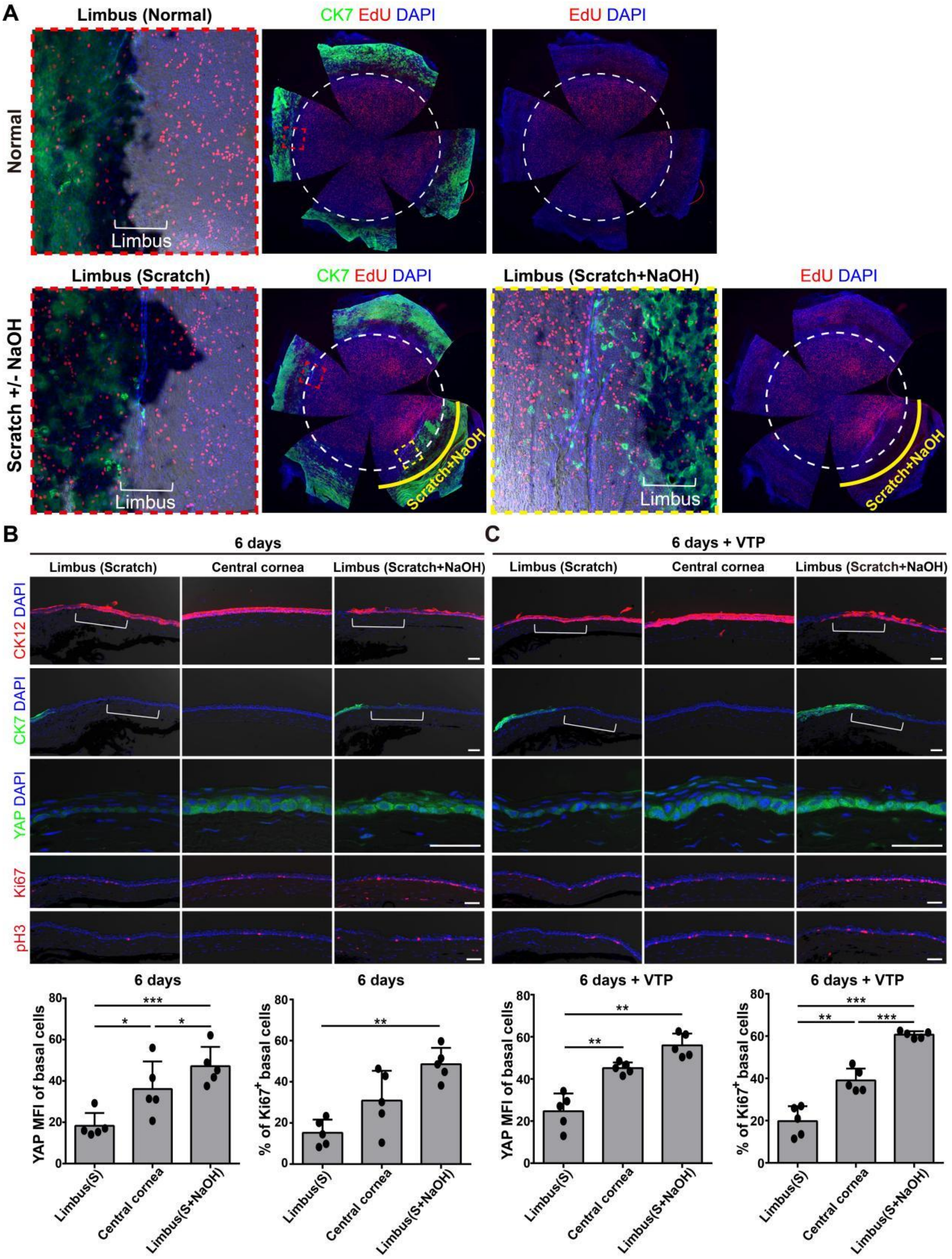


**Figure S7: The de-differentiation of CECs required an intact limbal niche and the suppression of YAP signal.**

(**A**) The whole-mount normal cornea and cornea of scratched limbus either with or without NaOH application were stained by CK7 and EdU. Dashed circles indicate the limbus. Note that the limbus of normal cornea contains a zone with low cell proliferation. The limbus where only scratch was applied recovered to the state of low cell proliferation, while limbus where both scratch and NaOH were applied maintained a state of high cell proliferation. (**B**) The frozen-section CK12, CK7, YAP, Ki67 and p-Histone H3 (pH3) immunostaining against cornea of scratched limbus either with or without NaOH application at 6 days. The MFI of YAP and the percentage of $Ki67^{+}$ cells in limbal basal cells were quantified. (**C**) The limbus was scratched either with or without NaOH application, and VTP was administrated for 4 days. The frozen-section CK12, CK7, YAP, Ki67 and pH3 immunostaining was performed at 6 days. The MFI of YAP and the percentage of $Ki67^+$ cells in limbal basal cells were quantified. Data are the mean ± SD, n=5 biological replicates; statistical analysis were performed by paired one-way ANOVA with Tukey's test (**B**, **C**). Scale bars, 50 um.

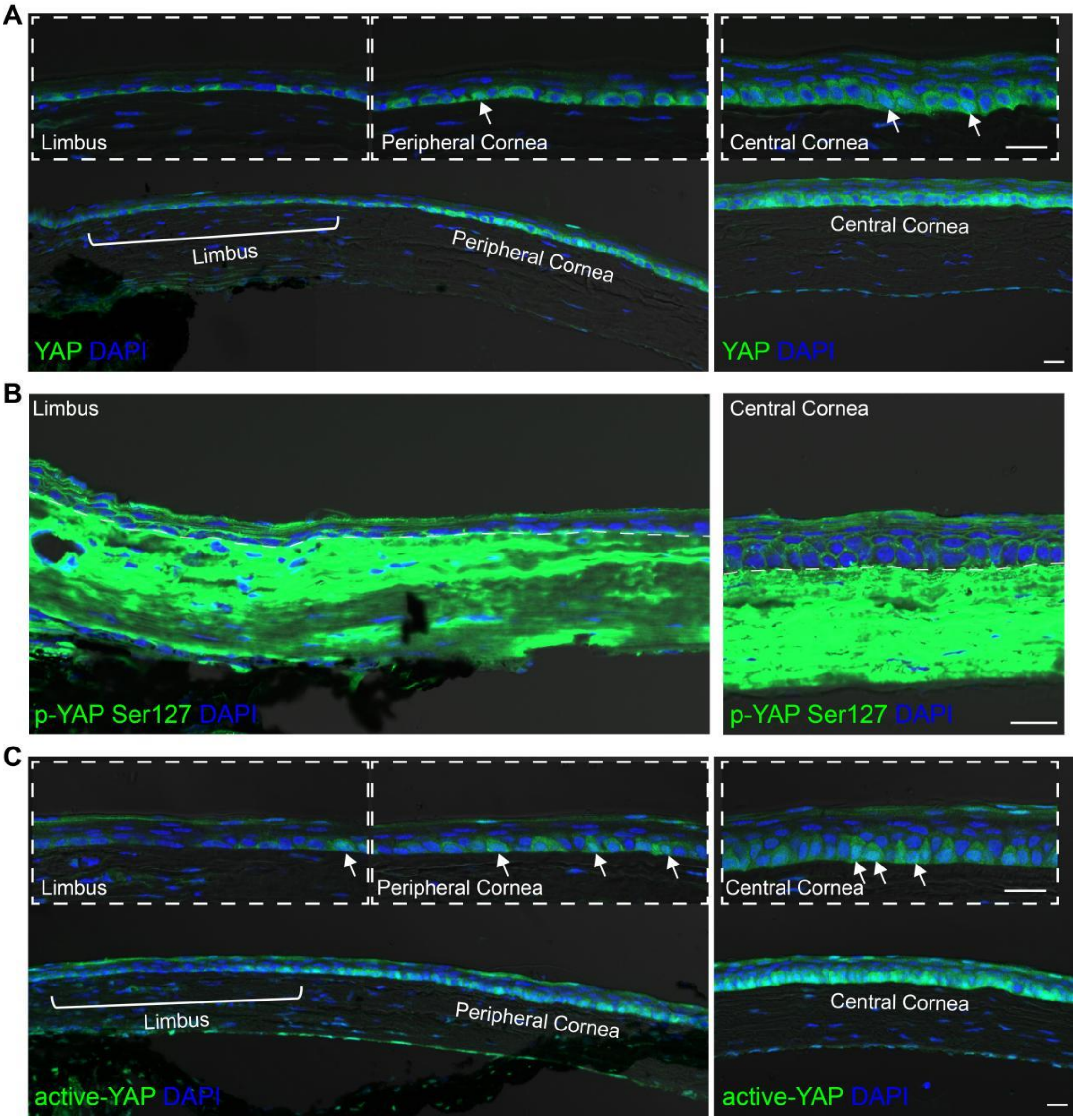


**Figure S8: The expression of YAP in cornea during normal homeostasis.**

(**A**) The expression of YAP at limbus, peripheral cornea and central cornea of mice. Note the higher fluorescence intensity at peripheral and central cornea than limbus. (**B**) The expression of inactive p-YAP (Ser127) at limbus, peripheral cornea and central cornea. (**C**) The expression of active-YAP (non-phosphorylated YAP) at limbus, peripheral cornea and central cornea. Note the higher fluorescence intensity at peripheral and central cornea than limbus. The arrows point CECs with nuclear YAP. Scale bars, 20 um.

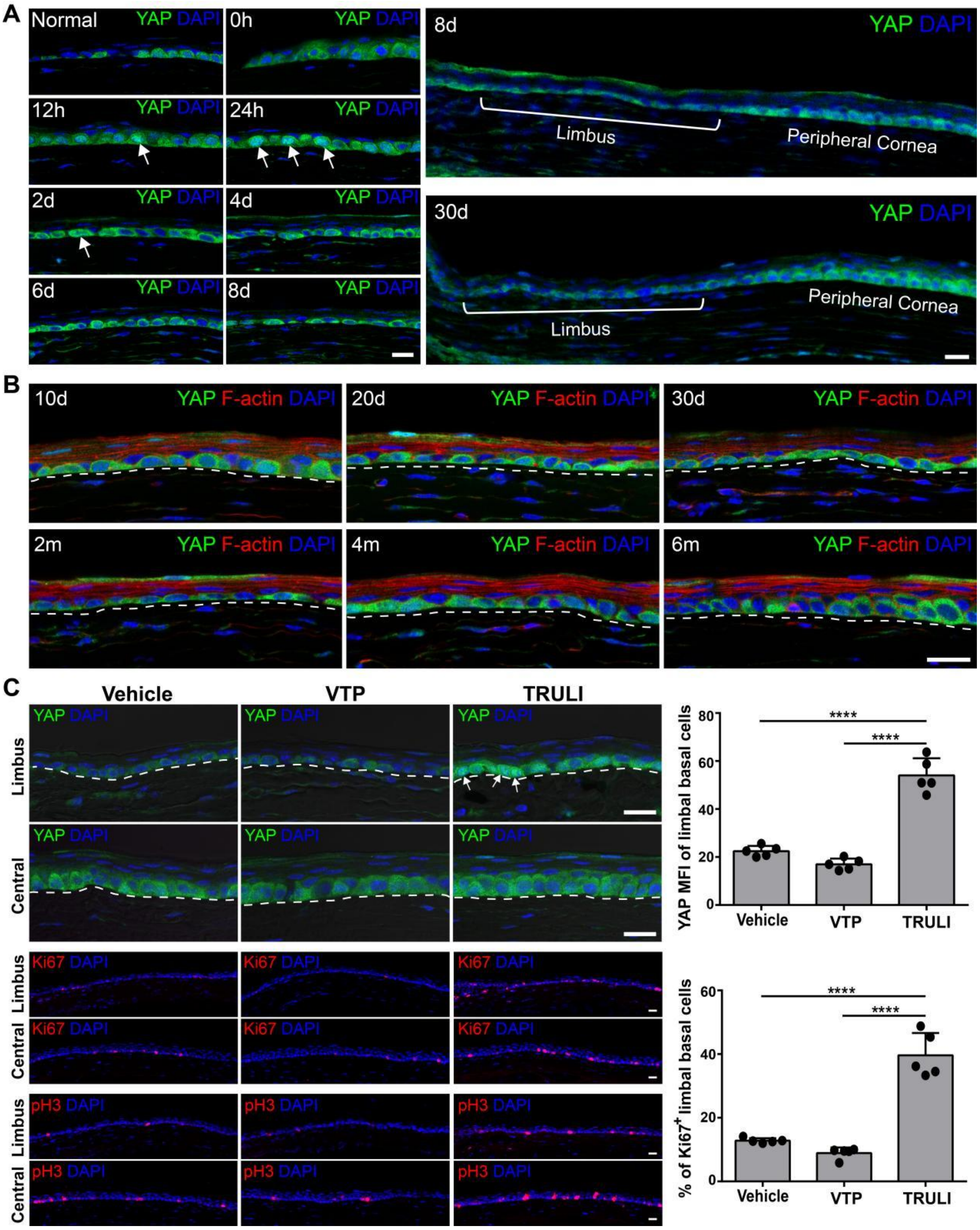


**Figure S9: The expression of YAP at limbus after the ablation of innate LESCs.**

(**A**, **B**) The expression of YAP at limbus at indicated days after the ablation of innate LESCs. (**C**) The expression of YAP, Ki67 and pH3 at limbus and central cornea at 6 days after the ablation of innate LESCs with the administration of vehicle or VTP or TRULI. Arrows point to CECs with nuclear YAP. The MFI of YAP and the percentage of $Ki67^{+}$ cells in limbal basal cells were quantified. Data are the mean ± SD, n=5 biological replicates; statistical analysis were performed by unpaired one-way ANOVA with Tukey's test. Scale bars, 20 um.

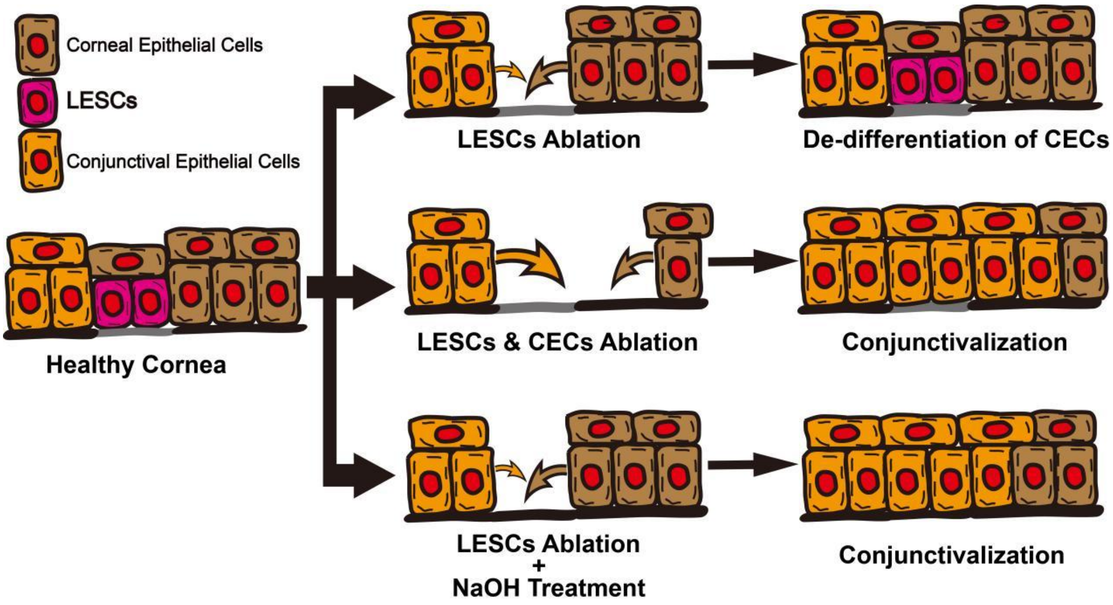


**Figure S10: The model of the de-differentiation of CECs.**

During normal homeostasis of cornea, LESCs at the limbus proliferate and differentiate into CECs to maintain the continuous renewal of corneal epithelium. After the ablation of innate LESCs, CECs migrate back to the limbus and de-differentiate into LESCs, thus maintaining the normal functions of corneal epithelium. When LESCs and CECs are ablated, conjunctival epithelial cells win the competition between CECs for the limbus, thus CECs can't occupy the limbus and de-differentiate into LESCs without the limbal niche, resulting in the conjunctivazation of cornea. In addition, when the limbal niche is destroyed by NaOH, although CECs can cover the limbus, they can't de-differentiate into LESCs and cornea become conjunctivazation.
